# rTg(Tau_P301L_)4510 mice exhibit increased VGLUT1 in hippocampal presynaptic glutamatergic vesicles and increased extracellular glutamate release

**DOI:** 10.1101/2022.04.21.489047

**Authors:** Erika Taipala, Jeremiah C. Pfitzer, Morgan Hellums, Miranda Reed, Michael W. Gramlich

## Abstract

The molecular pathways that contribute to the onset of symptoms in tauopathy models, including Alzheimer’s Disease (AD), are difficult to distinguish because multiple changes can happen simultaneously at different stages of disease progression. Understanding early synaptic alterations and their supporting molecular pathways is essential in order to develop better pharmacological targets to treat AD. Here we focus on an early onset rTg(Tau_P301L_)4510 tauopathy mouse model that exhibits hyperexcitability in hippocampal neurons of adult mice that is correlated with presynaptic changes and increased extracellular glutamate levels. However, it is not clear if increased extracellular glutamate is caused by presynaptic changes alone, or if presynaptic changes are a contributing factor among other factors. To determine whether pathogenic tau alters presynaptic function and glutamate release, we studied cultured hippocampal neurons at 14-18 DIV from animals of both sexes to measure presynaptic changes in tau_P301L_ positive mice. We observed that presynaptic vesicles exhibit increased vesicular glutamate transporter 1 (VGLUT1) using immunohistochemistry of fixed cells and an established pH-sensitive green fluorescent protein approach. We show that tau_P301L_ positive neurons exhibit a 40% increase in VGLUT1 per vesicle compared to tau_P301L_ negative littermates. Further, we use the extracellular glutamate reporter iGluSnFR to show that increased VGLUT1 per vesicle directly translates into a 40% increase in extracellular glutamate. Together, these results show that increased extracellular glutamate levels observed in tau_P301L_ mice are not caused by increased vesicle exocytosis probability but rather are directly related to increased VGLUT1 transporters per synaptic vesicle.

## 1 Introduction

Pathological hyperphosphorylation and aggregation of tau is a hallmark of neurodegenerative conditions known as tauopathies, including Alzheimer’s disease (AD), Parkinson’s disease (PD), Pick’s disease, and Frontotemporal dementia with Parkinsonson-17 (FTDP-17). While much of the work has focused on the role of pathological tau in disrupting postsynaptic signaling, there is growing interest in the role that pathological tau plays in presynaptic dysfunction (see (Wu et al., 2021) for review). Using the rTg(TauP301L)4510 mouse model (hereafter called tau_P301L_ pos), we have previously demonstrated *in vivo* that P301L tau expression increases glutamate release in the hippocampus, which correlated with memory deficits, at a time when tau pathology was subtle and before readily detectable neuron loss (Hunsberger et al., 2014). This increase in glutamate release was concomitant with a 40% increase in hippocampal levels of the vesicular glutamate transporter VGLUT1. Importantly, neither cognitive deficits nor the increase in glutamate release and VGLUT1 levels were observed in the rTg(TauWT)21221 mouse model that expresses wild-type 4R0N human tau at concentrations equivalent to P301L human tau in tau_P301L_ pos mice, but without the P301L mutation and associated tau pathology (Hunsberger et al., 2014). This suggests that overexpression of tau per se did not mediate these alterations, which we have also previously demonstrated for postsynaptic alterations associated with P301L tau (Hoover et al., 2010). Moreover, reducing VGLUT1 levels and glutamate release rescued cognitive deficits in tau_P301L_ pos mice (Hunsberger et al., 2015), suggesting the increased VGLUT1 and/or glutamate release may represent potential therapeutic targets for the treatment of early-stage tauopathies.

Prior studies have also shown VGLUT1 expression to be increased in the PS19 tauopathy mouse model in an age-dependent manner (Crescenzi et al., 2017), with levels increasing then decreasing with age. Likewise, while VGLUT1 is decreased in AD patients, particularly in the later stages (Kashani et al., 2008) patients with early stages of mild cognitive impairment (MCI) demonstrate increased VGLUT (Bell et al., 2007). Together, these results suggest that the changes in VGLUT1 expression may be age- or stage-dependent. This early increase in VGLUT1 in MCI patients may help explain the hippocampal hyperexcitability that is predictive of the degree and rate of cognitive decline, as well as the conversion to AD, that has been previously observed in MCI patients ((Mackenzie & Miller, 1994) and (see (Toniolo et al., 2020) for review).

A major limitation to understanding the mechanisms of tau-mediated changes in presynaptic transmission has been the ability to separately distinguish mechanisms at the single vesicle level in a physiological approach. Prior i*n vitro* cell cultures used immortalized cell lines or transfection of human tau (htau) in primary hippocampal cells (McInnes et al., 2018; Siano et al., 2019; Zhou et al., 2017), and have shown that tau mediates both an increase in VGLUT1 expression and a reduction in synaptic transmission. *In vivo* approaches have also been able to measure increased VGLUT1 expression and tau mediated reduction in synaptic transmission (Crescenzi et al., 2017), but could not directly measure the presynaptic contribution from the post-synaptic response. Separating the consequences of increased VGLUT1 from the decrease in synaptic transmission is essential because changes in VGLUT1 expression have been shown to increase glutamate concentration per vesicle (Wilson et al., 2005) and changes in release probability (Herman et al., 2014). Thus, tau mediated decrease in exocytosis and tau mediated increase in VGLUT1 expression levels can have counteracting effects on presynaptic transmission. These limitations have prevented a complete mechanistic understanding of the effect of tau-mediated alterations in VGLUT1 levels on presynaptic transmission, which we address here.

Another limitation is separating the role of tau-mediated changes in frequency-dependent synaptic transmission. Normal synaptic transmission utilizes a complex train of stimulus from low frequencies (1 Hz) to high frequencies (100 Hz) (Abbott & Regehr, 2004; Klyachko & Stevens, 2006). Multiple studies of tauopathy models have shown hyperexcitable periods of higher frequency activity (> 20Hz) occurring for sustained bouts (Kazim et al., 2017). It is not clear if these periods of hyperexcitability are mediated by changes in a frequency-dependent alteration in presynaptic transmission. If tau mediates frequency-dependent release of glutamate, then this represents an important potential pathway of disease progression, because prior studies have demonstrated a glutamate-mediated exocytosis of tau as a potential mechanism for the trans-synaptic spread of tau pathology (Liu et al., 2012; Pooler et al., 2013; Yamada et al., 2014), and neuronal hyperexcitability represents one potential source for the pathogenesis of tauopathies (see (Toniolo et al., 2020) for review). However, no careful study of frequency-dependent presynaptic glutamate release has been explored in cell culture models.

In the present study, we use nanometer fluorescence microscopy and computational analysis techniques to quantitatively explore how presynaptic vesicle VGLUT1 is altered in a P301L tauopathy mouse model, as well as the resulting consequences on glutamate release in postnatal day (PND5) primary hippocampal cell cultures of neurons grown on astrocytes. Presynaptic alterations were measured at 14-18 days in vitro (DIV). We isolate presynaptic transmission from postsynaptic response using established postsynaptic AMPA and NMDA receptor blockers during experiments (Gramlich & Klyachko, 2017; Maschi et al., 2018; Maschi & Klyachko, 2017; Voglmaier et al., 2006). We compare tau_P301L_ pos neurons grown on tau_P301L_ pos astrocytes to tau_P301L_ neg neurons grown on tau_P301L_ neg astrocytes at the same time-point. We directly image single vesicle VGLUT1 levels using the pH-sensitive GFP fluorescent protein pHlourin (Maschi & Klyachko, 2017; Voglmaier et al., 2006). Further, we image extracellular glutamate levels using the GFP fluorescent plasma membrane reporter iGluSnFR (Marvin et al., 2018).

We observed an overall increase in pHluorin-VGLUT1 intensity during a single bout of stimulation in tau_P301L_ pos neurons compared to tau_P301L_ neg neurons. The increased pHluorin-VGLUT1 intensity was independent of stimulation frequency, or single release probability per stimulation. We observed a 40% increase in pHluorin-VGLUT1 intensity per vesicle in tau_P301L_ pos neurons compared to tau_P301L_ neg neurons during stimulation. We observed that the increase VGLUT1 per vesicle correlates with a 40% increase in iGluSnFR glutamate reporter intensity, which supports the hypothesis that increased VGLUT1 transporters per vesicle results in extracellular glutamate released during stimulation. Finally, we use our results to estimate the number of VGLUT1 transporters per vesicle, the number of vesicles released during stimulation, and the number of extracellular glutamate molecules in the synaptic cleft during stimulation.

## 2 Results

### 2.1 Tau_P301L_ positive neurons exhibit increased pHluorin-VGLUT1 fluorescence intensity compared to tau_P301L_ negative neurons

To determine if previously observed increased VGLUT1 expression (Hunsberger et al., 2014) exists in tau_P301L_ pos hippocampal neurons compared to their transgene negative littermates (hereafter called tau_P301L_ neg), we used the established pH-sensitive fluorescent protein-based indicator pHluorin-VGLUT1 approach in our cultured cells (Maschi et al., 2018; Maschi & Klyachko, 2017; Voglmaier et al., 2006). We fluorescently imaged transfected cells before, during, and after a single bout of electrical stimulation, which results in an increase in pHluorin intensity due to exposure of synaptic vesicles to the neutral pH of the extracellular environment (Fig. 1A). The resulting observed pHluorin-VGLUT1 intensity is then quantized for each vesicle released (ΔF/pulse, Fig. 1A).

**Figure 1:**
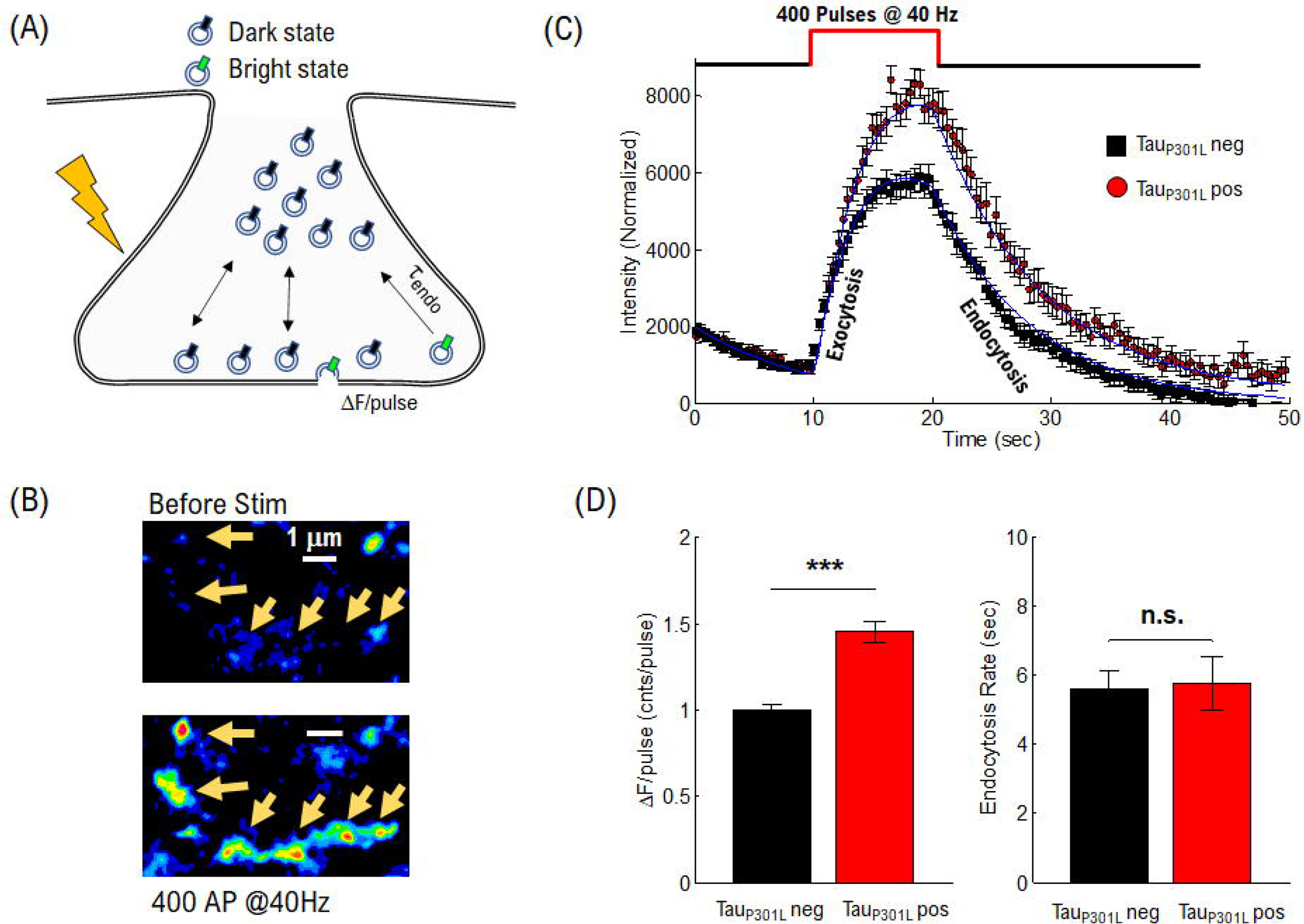
Tau_P301L_ pos neurons exhibit increased overall pHluorin-VGLUT1 intensity compared to tau_P301L_ neg neurons. (A) Representation of pHluorin-VGLUT1 intensity observed during presynaptic transmission (B) Example of raw pHluorin-VGLUT1 data observed before and directly after a single bout of 40 Hz stimulation (C) Comparison of pHluorin-VGLUT1 intensity before, during, and after 40 Hz stimulation for tau_P301L_ neg (Black) and tau_P301L_ pos (red) neurons (D) Comparison of intensity/pulse and endocytosis rate for tau_P301L_ neg (Black) and tau_P301L_ pos (red) neurons. tau_P301L_ neg N = 257 presynapses; tau_P301L_ pos N = 160 presynapses; *** = P < 0.001 for Mann-Whitney U-test. NS=not

Specifically, we chose a stimulation frequency of 40Hz in order to drive a significant portion of the recycling pool to release in order to observe a difference in pHluorin-VGLUT1 intensity in tau_P301L_ pos neurons as compared to tau_P301L_ neg controls. Natural spike trains in hippocampal neurons range from 1 Hz up to ∼100 Hz (Klyachko & Stevens, 2006). However, hippocampal neurons exhibit an increase in spiking rates characterized by extended bouts (∼10 sec) of higher-frequency spiking rates (>=20 Hz) prior to neurodegeneration in AD models (Kazim et al., 2017). Consequently, a single 10 sec bout of electrical stimulation at a constant rate of 40 Hz will model this hyperexcitable state and allow for a clear observable difference in pHluorin-VGLUT1 intensity (Fig. 1B).

Measurably significant differences occurred in the quantitative pHluorin-VGLUT1 intensity observed in tau_P301L_ pos neurons as compared to tau_P301L_ neg controls. The observed pHluorin-VGLUT1 intensity exhibited an initial increase with stimulation (exocytosis, Fig. 1C) followed by a saturation, and then a reduction in pHluorin-VGLUT1 intensity (endocytosis, Fig. 1C). This is consistent with every previous pHluorin-VGLUT1 study in hippocampal neurons (Maschi et al., 2018; Maschi & Klyachko, 2017; Voglmaier et al., 2006). However, in this present study, tau_P301L_ pos (Red squares, Fig. 1C, N = 160) exhibited an overall increase in pHluorin-VGLUT1 intensity as compared to tau_P301L_ neg controls (Black circles, Fig. 1C, N = 257).

There are two potential contributions from vesicle recycling mechanics that could result in the differences between tau_P301L_ pos and tau_P301L_ neg intensity. Because each exocytosed vesicle immediately begins the endocytosis process, observed pHluorin-VGLUT1 intensity simultaneously includes increasing intensity from exocytosis (ΔF/pulse, Fig. 1A) and decreasing intensity from endocytosis (τ_endo_, Fig. 1A) (Chanaday & Kavalali, 2018). To delineate their relative contributions, we developed a continuum fit function that can distinguish exocytosis and endocytosis parameters, while reproducing observed intensities:

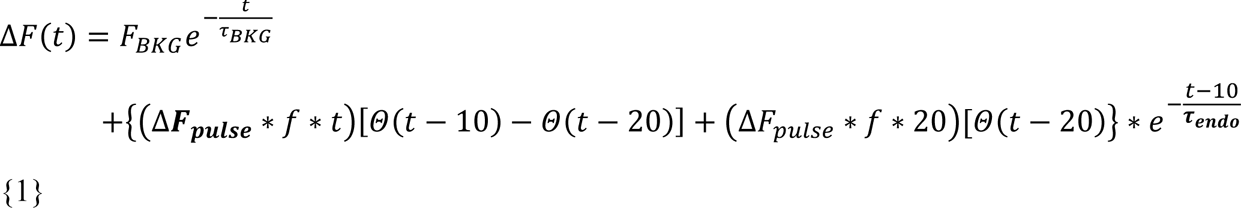

where F_BKG_ is the starting background intensity, τ_BKG_ is the background photobleaching rate constant, ΔF_pulse_ is the pHluorin intensity increase per pulse, f is the stimulation frequency, Θ is the Heaviside function, τ_endo_ is the endocytosis rate constant.

This continuum function reproduced the observed pHluorin-VGLUT1 intensity with time for both tau_P301L_ pos and tau_P301L_ neg neurons (Blue Solid Lines, Fig. 1C). From this analysis, we were able to separately quantify the intensity increase per stimulus (ΔF_pulse_, Left Panel, Fig. 1D) and the endocytosis rate (τ_endo_, Right Panel, Fig. 1D). We normalized the tau_P301L_ pos values to the tau_P301L_ neg control to highlight differences between conditions and found that the ΔF_pulse_ for tau_P301L_ pos neurons showed an increase of 1.4x the tau_P301L_ neg controls (1.45 ± 0.07, P=1.5E-8; Mann-Whitney U-test comparisons), whereas the endocytosis rate was the same for both tau_P301L_ pos and tau_P301L_ neg neurons (tau_P301L_ neg:5.57 ± 0.5; tau_P301L_ pos: 5.76 ± 0.8;P > 0.4, Mann-Whitney U-test). Further, the endocytosis rate is consistent with previously observed endocytosis rates in hippocampal neurons using pHluorin-VGLUT1 intensity analysis (Chanaday & Kavalali, 2018).

The increase in intensity per stimulus, but not endocytosis rate, in tau_P301L_ pos neurons suggests that a change in either (a) exocytosis mechanics or (b) the amount of VGLUT1 per vesicle or both. We will focus the remainder of the present study to distinguish if either (a), (b), or both occur in tau_P301L_ pos neurons.

### 2.2 Computational model of presynaptic exocytosis distinguishes pathways to increased tau_P301L_ pos pHluorin-VGLUT1 intensity

To understand how tau_P301L_ pos neurons could exhibit increased pHluorin-VGLUT1 intensity compared to tau_P301L_ neg neurons, we developed a computational simulation model of the experimental results (Fig. 2). Our model is based on the established binomial model of presynaptic transmission (Lanore & Silver, 2016; Reid & Clements, 1999; Scheuss & Neher, 2001). The established model predicts a presynaptic response to a single stimulus as:

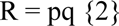

where R is the response, p is the probability of release per stimulus, and q is the post-synaptic response.

**Figure 2:**
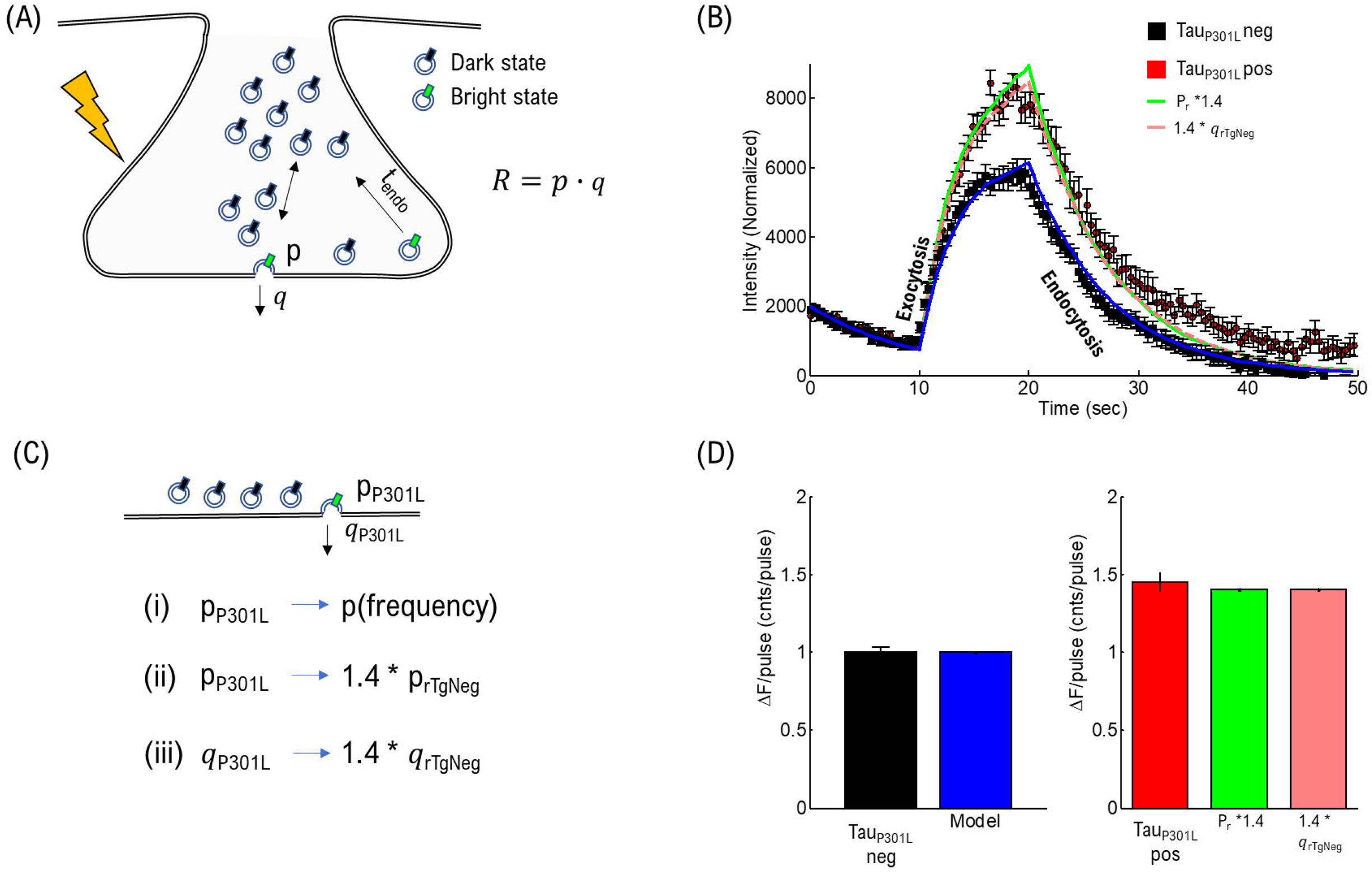
Computational model of presynaptic exocytosis. (A) Cartoon representation of computational model of presynaptic release in tau_P301L_ neg and tau_P301L_ pos neurons (B) Comparison of pHluorin data and computational model (C) Hypothesized pathways for tau_P301L_ pos pHluorin-VGLUT1 intensity increase (D) Quantitative comparison of resulting ΔF/pulse for experimental results and computational models

Here we modify the traditional model to reproduce pHluorin-VGLUT1 intensity as (See Supplemental Methods):

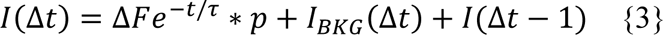

where each simulation time-step (Δt) includes the intensity from the previous simulation time-step, a total background intensity (I_BKG_, equivalent to eqn. 1), and a probabilistic increase (p) in intensity given by a per vesicle count (ΔF, equivalent to eqn. 1) from each stimulus pulse, and each vesicle has the same endocytosis rate (τ, equivalent to eqn. 1).

In the present study, we used parameters for endocytosis and release probability based on experimentally determined values obtained in this study. We required that all vesicles have the same endocytosis rate (τ = 7 sec^-1^), found from fits in eqn. 1. To model tau_P301L_ neg single vesicle release probability per stimulus pulse, we used the experimentally observed release probability (P = 0.1) found in the present study (Fig. 4D). We allowed the intensity per vesicle and overall release probability per site to change in order to model observed tau_P301L_ pos intensity.

To confirm the validity of this model, we first compared its results to observed tau_P301L_ neg pHluorin-VGLUT1 intensity. This model reproduced observed experimental tau_P301L_ neg pHluorin-VGLUT1 intensity (Solid line through tau_P301L_ neg, Fig. 2B) with equivalently good agreement as the continuum model in eqn. 1 (Fig. 1 C). Further, this model results in the same quantitative intensity per pulse as the experimentally observed (Left Panel Fig. 2 D). Thus, this model provides a sufficient representation of observed presynaptic release for modelling tau_P301L_ pos pHluorin-VGLUT1.

We then proposed three possible pathways by which tau_P301L_ pos pHluorin-VGLUT1 intensity could be greater than tau_P301L_ neg pHluorin-VGLUT1 intensity (Fig. 2C):

(i) tau_P301L_ pos presynapses exhibit a frequency-dependent increase (Section 2.3).
(ii) tau_P301L_ pos release sites exhibit an overall 1.4x increase in release-probability (Section 2.4).
(iii) tau_P301L_ pos vesicles exhibit 1.4x pHluorin-VGLUT1 intensity per vesicle compared to tau_P301L_ neg (Section 2.5).

The first pathway is tested experimentally in the following section 2.3. The second pathway, modeled computationally as an increased p-value (p, eqn. 3), reproduces observed tau_P301L_ pos intensity (Solid Line, Fig. 2B); this pathway is experimentally tested in section 2.4. The third pathway, modeled as an increase in the pHluorin-VGLUT1 intensity per vesicle (ΔF, eqn. 3), results in the same observed tau_P301L_ pos pHluorin-VGLUT1 intensity (Dashed Line overlaps solid line, Fig. 2B); this pathway will be experimentally tested in section 2.5.

### 2.3 Tau_P301L_ pos neurons exhibit a frequency independent pHluorin-VGLUT1 intensity increase as compared tau_P301L_ neg neurons

Tau has been implicated in facilitating hyperexcitability in mouse models of AD as well as models of epilepsy (Ittner et al., 2010; Palop et al., 2007; Peters et al., 2020; Roberson et al., 2007, 2011), and it is possible that pHluorin-VGLUT1 intensity in tau_P301L_ pos mice may change as a function of neuronal stimulation.

To establish if any stimulation frequency-dependent increase in vesicle release events exists in tau_P301L_ pos hippocampal neurons, we performed pHluorin-VGLUT1 intensity experiments as a function stimulus frequency using frequencies that range from moderate (10 Hz) to potential hyperexcitable frequencies (40 Hz) (Fig. 3). For a consistent comparison of pHluorin-VGLUT1 intensity across stimulation frequencies, we constrained the total time of stimulation (10 sec) by limiting the total number of pulses. We then compared the observed increase in pHluorin-VGLUT1 intensity during stimulation for both tau_P301L_ pos and tau_P301L_ neg neurons (10 Hz to 40 Hz). Lastly, we separated the baseline and stimulation time (Fig. 3 A,B) from the post-stimulation pHluorin-VGLUT1 intensity decay (Fig. 3 D,E) to separately compare exocytosis and endocytosis processes.

**Figure 3:**
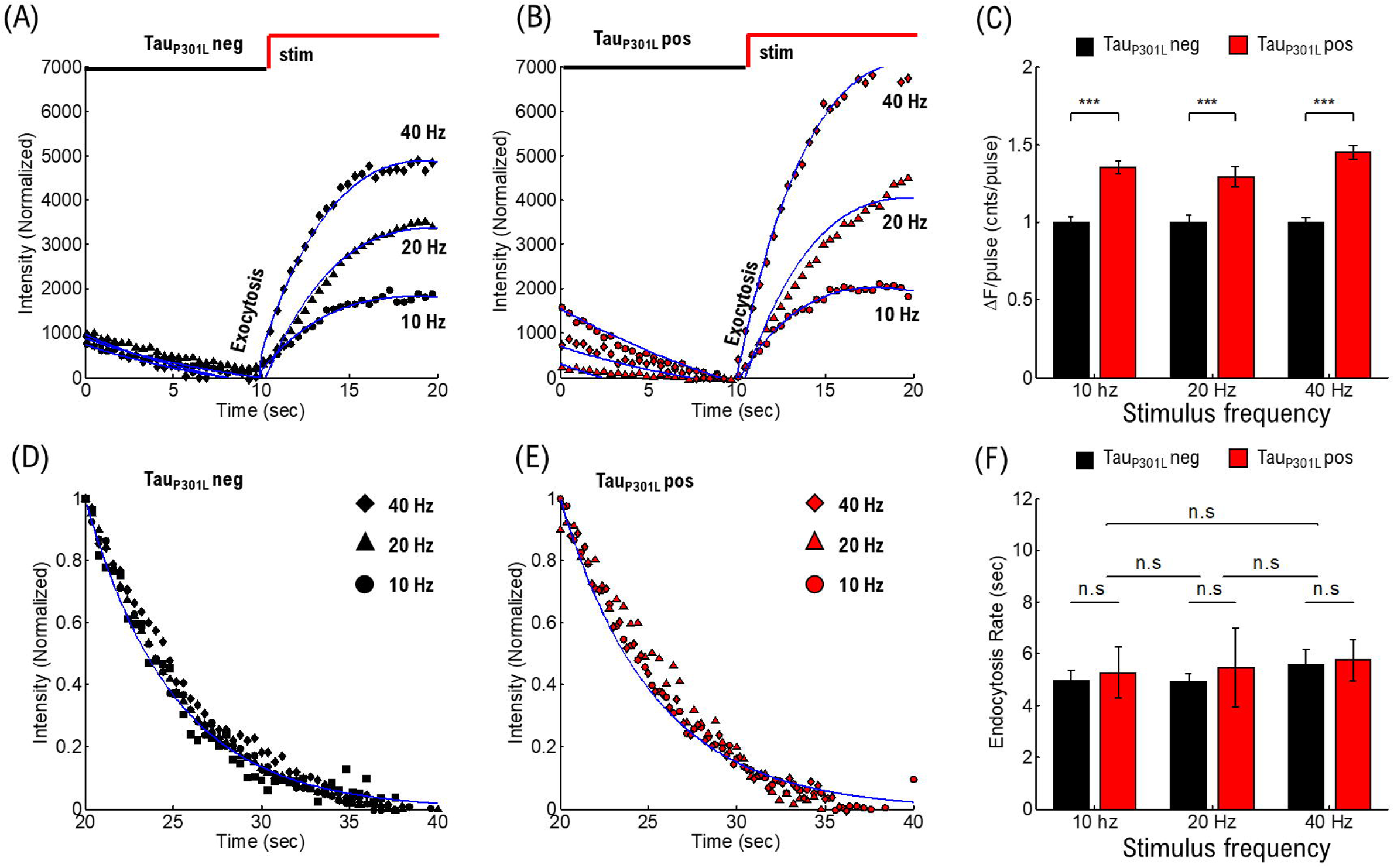
Frequency-dependent stimulation does not contribute to increased tau_P301L_ pos pHluorin-VGLUT intensity. (A) Comparison of pHluorin-VGLUT1 intensity before (0-10 sec) and during (10-20 sec) stimulation as a function of stimulation frequency for tau_P301L_ neg neurons. (B) Comparison of pHluorin-VGLUT1 intensity before (0-10 sec) and during (10-20 sec) stimulation as a function of stimulation frequency for tau_P301L_ pos neurons. (C) ΔF/pulse parameter from eqn. 1 fit to data (Blue Lines in A,B). Values at each stimulation frequency are normalized to the tau_P301L_ neg value for that frequency. (D) Comparison of pHluorin-VGLUT1 intensity decay after stimulation (20-40 sec) as a function of stimulation frequency for tau_P301L_ neg neurons. (E) Comparison of pHluorin-VGLUT1intensity decay after stimulation (20-40 sec) as a function of stimulation frequency for tau_P301L_ pos neurons. (F) Endocytosis Rate from exponential decay fits (Blue Line in D,E). Each curve fit separately. Errors are twice the 95% confidence interval value of fits to data. *** = P<0.01; Statistical results from Mann-Whitney U.

To quantify the difference between tau_P301L_ neg and tau_P301L_ pos intensity as a function of stimulation frequency, we fit the frequency-dependent curves to our continuum model (eqn. 1, Blue Lines in Fig. 3 A,B). Each curve was fit with the same constraints on endocytosis rate and background intensity, resulting in a single fit parameter corresponding to the difference in single vesicle intensity per pulse (ΔF/pulse). Lastly, to directly compare changes as a function of frequency, we normalized the ΔF/pulse to the tau_P301L_ neg parameter (Fig. 3C). We found that tau_P301L_ pos ΔF/pulse was consistently higher for tau_P301L_ pos for all stimulation conditions (10 Hz: 1.35 ± 0.05, P=7.5E-4; 20 Hz: 1.3 ± 0.1, P=2.9E-4; 40 Hz: 1.45 ± 0.07, P=1.5E-8; Mann-Whitney U-test comparisons), suggesting that the increased pHluorin-VGLUT1 intensity observed in tau_P301L_ pos neurons is not frequency-dependent.

To further support that tau_P301L_ pos pHluorin-VGLUT1 intensity increases are independent of stimulation frequency, we next established if the endocytosis rate is dependent upon stimulation frequency. We compared the endocytosis rate for all stimulation frequencies by normalizing by background and peak intensity. If the endocytosis rate is independent of stimulation frequency, then the normalized curves should all exhibit the same decay rate.

We observed that both tau_P301L_ neg (Fig. 3D) and tau_P301L_ pos (Fig. 3E) decay curves exhibit the same normalized decay rate as a function of time after stimulus. All curves were fit separately to a single exponential decay function with a single example curve shown for each condition (Blue Line). The resulting fit rate-constant was consistent for both tau_P301L_ neg (10 Hz: 4.9 ± 0.4; 20 Hz: 5.3 ± 0.5; 40 Hz: 5.6 ± 0.5; P > 0.4 for all curves, Mann-Whitney U-test comparisons) and tau_P301L_ pos neurons (10 Hz: 5.3 ± 0.5; 20 Hz: 5.1 ± 1.5; 40 Hz: 5.8 ± 0.8; P > 0.4 for all curves, Mann-Whitney U-test comparisons). Further, there was no statistical difference between any of the stimulation frequency results (P > 0.35 for all tau_P301L_ neg/tau_P301L_ pos comparisons, Mann-Whitney U-test comparisons).

These combined exocytosis and endocytosis results show that observed increases in tau_P301L_ pos pHluorin-VGLUT1 intensity as compared to tau_P301L_ neg controls is not a frequency dependent release effect. Further, the observed decay rates are consistent with previously observed endocytosis rates using the same pHluorin-VGLUT1 approach (Chanaday & Kavalali, 2018; Voglmaier et al., 2006). However, the frequency-dependent results *do not* discount the possibility that tau_P301L_ pos neurons exhibit an overall increase in release probability hypothesized by the computational simulation (Fig. 2) independent of stimulation frequency.

### 2.4 Tau_P301L_ pos neurons exhibit the same release-probability as tau_P301L_ neg neurons

We next focused on the hypothesis that single vesicle release probability is increased in tau_P301L_ pos neurons compared to tau_P301L_ neg controls (pathway (ii)), by using single vesicle release events at low stimulation frequencies to establish if overall tau_P301L_ pos release probabilities were higher than tau_P301L_ neg controls (Fig. 3). Our approach is similar to a previously established method of observing single pHluorin-VGLUT1 intensity events at low stimulation frequencies (Maschi & Klyachko, 2017). Briefly, we stimulate cultured neurons at a fixed frequency (1 Hz, 2 Hz, or 10 Hz) for 30 seconds and observe pHluorin-VGLUT1 intensity during stimulation. We count the number of pulses until the *first release event* is observed (Fig. 4 A,B). For low stimulation frequencies (1 or 2 Hz), a single release event was typically observed for several frames before endocytosis occurred (Fig. 4A); whereas, for a moderate stimulation frequency (10 Hz), recurring events resulted in increasing intensity after the initial release (Fig. 4B). Consequently, we take the pulse just before the onset of intensity as the first release event (Green Arrow, Fig. 4A). We then aggregate the number of pulses before the first release event is observed for each condition to determine the release probability (Red arrows, Fig. 4AB, and Fig. 4C). Lastly, we then define the release probability as the inverse of the mean number of pulses until release (Fig. 4A):

**Figure 4:**
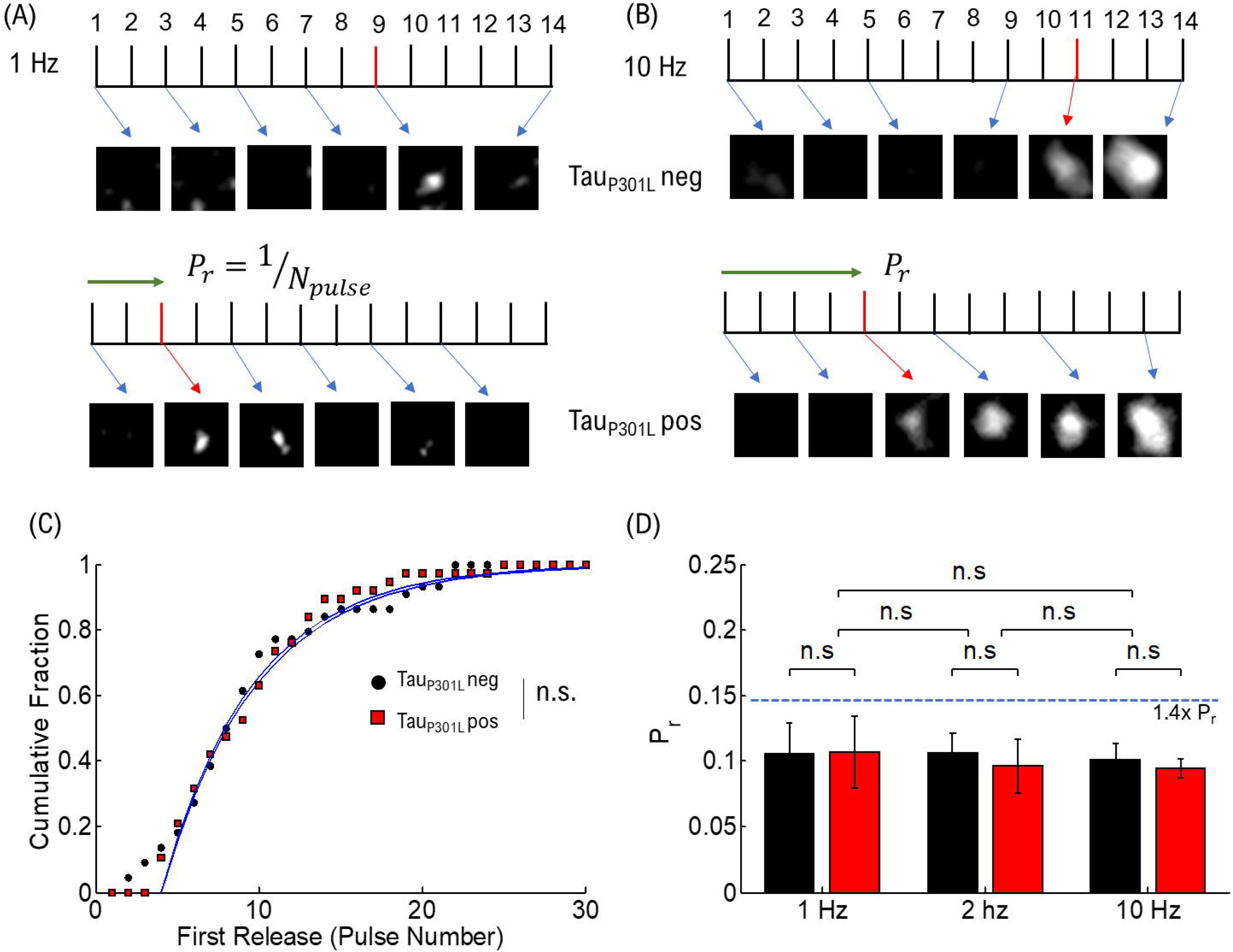
Release probability does not contribute to increases in tau_P301L_ pos pHluorin-VGLUT1 intensity. (A) Single Vesicle Release events as a function of Pulse Number for tau_P301L_ neg and tau_P301L_ pos conditions. (B) Initial Vesicle Release events as a function of Pulse Number for tau_P301L_ neg and tau_P301L_ pos conditions. (C) Aggregate cumulative distributions of number of pulses until the first release event is observed. (D) Release Probability as a function of stimulation frequency for tau_P301L_ neg (black) and tau_P301L_ pos conditions (red). All values reported as mean ± SEM. Statistical tests are two-tailed KS-test of cumulative fractions.

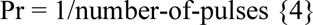

We note that the definition in our present study differs from previous pHluorin-VGLUT1 measurements, where a single pulse is initiated and a small number of release events are counted (Maschi et al., 2018; Maschi & Klyachko, 2017), and results in an elevated P_r_ due to residual calcium increasing the probability of release for each pulse. However, this approach presents an effective quantitative comparison between tau_P301L_ pos and tau_P301L_ neg vesicle release probability because they are being compared using the same metric.

Single release events were easily discerned for both tau_P301L_ pos and tau_P301L_ neg conditions (Fig. 4A). Likewise, initial release events were discerned for 10 Hz stimulation frequency (Fig. 4B). Aggregate cumulative fraction of release times for both tau_P301L_ pos (N_pulse_ = 9.33 ± 0.9) and tau_P301L_ neg (N_pulse_ = 9.45 ± 0.6) neurons showed the same pulse-to-first-release at 1 Hz stimulation frequency (P = 0.95, two-tailed KS-test, Fig. 4C). We performed the same statistical analysis for 2 Hz and 10 Hz. We then calculated the release probability as a function of stimulation for 1Hz (tau_P301L_ neg: 0.107 ± 0.009, tau_P301L_ pos: 0.109 ± 0.007; P = 0.95, two-tailed KS-test, Fig. 4D), 2 Hz (tau_P301L_ neg: 0.106 ± 0.009, tau_P301L_ pos: 0.096 ± 0.007; P = 0.95, two-tailed KS-test, Fig. 4D), and 10 Hz (tau_P301L_ neg: 0.10 ± 0.009, tau_P301L_ pos: 0.095 ± 0.007; P = 0.95, two-tailed KS-test, Fig. 4D).

If tau_P301L_ pos vesicle release probability was higher by 40% tau_P301L_ neg as hypothesized in pathway (ii), then the measured tau_P301L_ pos P_r_ should be higher than measured tau_P301L_ neg values. Since all tau_P301L_ neg values were P_r_∼0.1, all observed tau_P301L_ pos values *should have been* ∼0.14 (Dotted Blue Line, Fig. 4D). None of the measured tau_P301L_ pos values were significantly higher than tau_P301L_ neg (Fig. 4D). Further, none of the resulting tau_P301L_ pos and tau_P301L_ neg vesicle release probabilities were statistically different from each other for the same frequencies or across frequencies. Consequently, there is no difference in vesicle release probability in tau_P301L_ pos neurons as compared to tau_P301L_ neg.

The vesicle release probability results show that observed increases in pHluorin-VGLUT1 intensity (Fig. 1, 3) is not caused by an overall increase in vesicle release probability in tau_P301L_ pos neurons as compared to tau_P301L_ neg controls. This suggests that proposed pathway (ii) is not the mechanism by which tau_P301L_ pos pHluorin-VGLUT1 intensity is larger than tau_P301L_ neg.

### 2.5 Tau_P301L_ pos neurons exhibit quantized increase in pHluorin-VGLUT1 intensity-per-vesicle as compared to tau_P301L_ neg neurons

We last sought to establish that increased pHluorin-VGLUT1 intensity in tau_P301L_ pos neurons is caused exclusively by increased pHluorin-VGLUT1 intensity per vesicle by distinguishing the pHluorin-VGLUT1 intensity per single vesicle release events (Fig. 5). If single vesicle pHluorin-VGLUT1 intensity alone reproduces observed differences between tau_P301L_ neg and tau_P301L_ pos neurons, then single vesicle release event intensities should be ∼1.4x the pHluorin-VGLUT1 intensity per tau_P301L_ neg vesicle.

**Figure 5:**
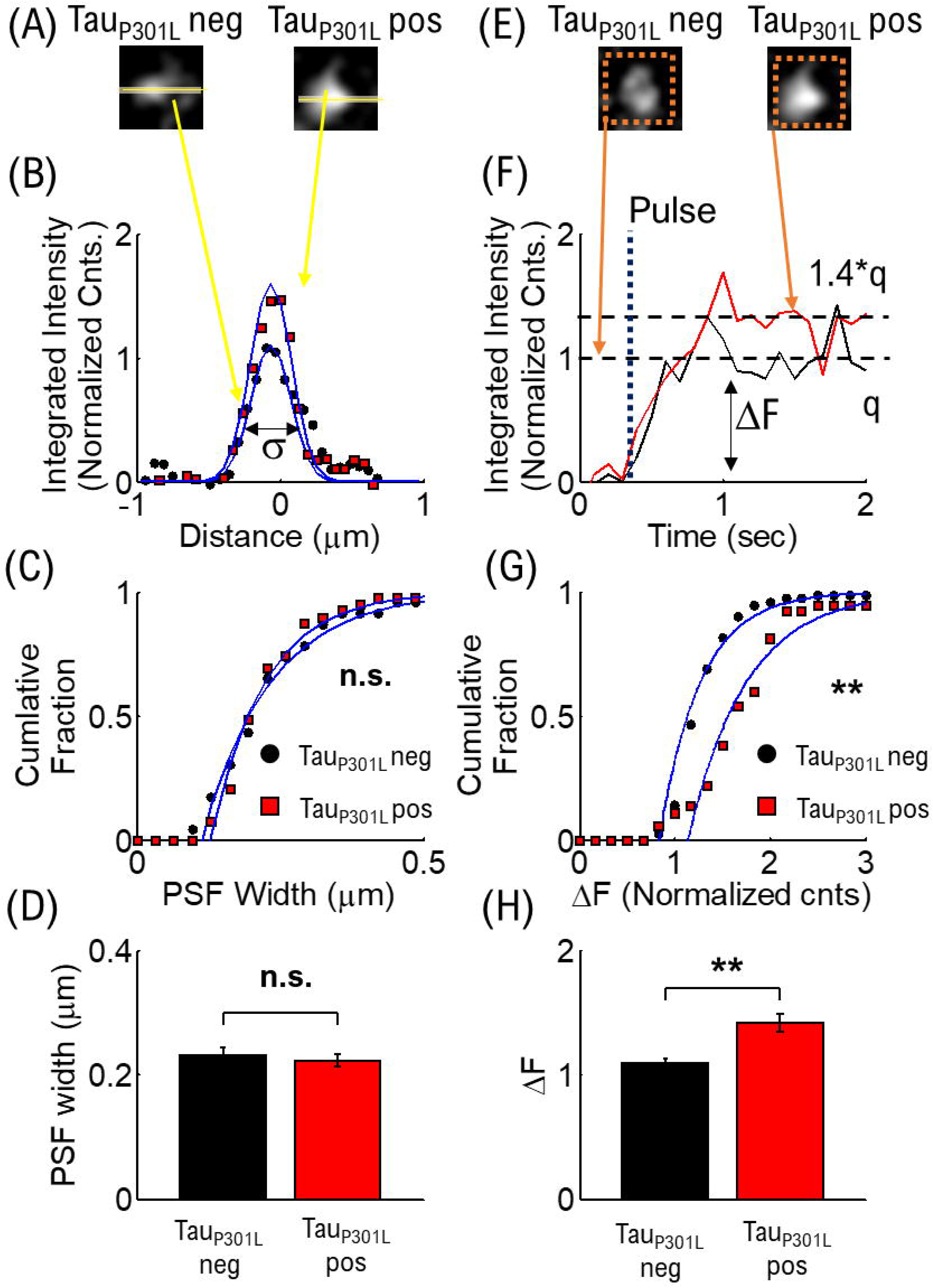
Quantized single vesicle pHlluorin- VGLUT1 intensity accounts for overall increased tau_P301L_ pos pHluorin intensity compared to tau_P301L_ neg neurons. (A) Example single vesicle events and line-profile analysis (Solid Yellow lines) for tau_P301L_ neg and tau_P301L_ pos neurons. (B) Example PSF intensity of tau_P301L_ neg (Black circles) and tau_P301L_ pos (Red squares) neurons from line-profiles in (A). Intensity has been background subtracted and normalized to tau_P301L_ neg peak intensity. Single gaussian fits to data (Blue solid lines), with PSF-width (σ) used to determine MVR. (C) Cumulative fraction distributions of tau_P301L_ neg (Black circles, N = 22) and tau_P301L_ pos (Red squares, N = 38) of single vesicle PSF-width (σ in B). Cumulative distributions fit to a single cumulative fit function (Blue Lines). (D) Mean values from cumulative fits to data (Blue lines in C), with errors from 95% confidence intervals of fits. (E) Example single vesicle events and integrated intensity analysis (Dashed Orange Boxes) for tau_P301L_ neg and tau_P301L_ pos neurons. (F) Sample integrated intensities traces for tau_P301L_ neg (Black line) and tau_P301L_ pos (Red Line) background subtracted and normalized to the average tau_P301L_ neg intensity. Single release events are observed after a stimulus pulse (Dotted Line labeled Pulse). Tau_P301L_ pos neurons exhibit larger intensity (ΔF) increase (1.4*q) than tauP301L neg (q). (G) Cumulative fraction of aggregate single vesicle intensity (ΔF) after stimulus relative to background. All data normalized to average tau_P301L_ neg intensity. Both curves fit with a cumulative fit function (Blue solid lines). (H) Mean ± SEM intensity per single vesicle release event for tau_P301L_ neg and tau_P301L_ pos from cumulative fractions in G. ** = P < 0.01, KS two-tailed test.

Before establishing intensity per vesicle, we sought to establish that only a single vesicle released during a stimulus event. Previous pHluorin-VGLUT1 experiments have established that presynapses can exhibit multiple vesicle release (MVR) (Maschi et al., 2018, 2021; Maschi & Klyachko, 2017), which may also increase observed pHluorin-VGLUT1 intensities. To establish that single vesicle releases are the predominant process in tau_P301L_ pos, we quantified the point-spread-function (PSF) per release event by obtaining line-profile counts for vesicles during the first observed frame (Fig. 5AB). We fit the PSF line profile to a gaussian function (Blue lines in Fig. 5B) to determine the width of each observed release event (σ, Fig. 5B). If multiple vesicles were released at the same time in tau_P301L_ pos as compared to tau_P301L_ neg, then the PSF-width increase (Maschi et al., 2021). We then aggregated the PSF of release events where only background intensity was observed before the release and where a single intensity step increase was observed within a 5-frame window (Fig. 5F) in order to prevent bias of asynchronous or spontaneous release events. These restrictions resulted in a limited but comparable number of events for both tau_P301L_ pos (N = 38) and tau_P301L_ neg conditions (N = 22), however we observed that the average width per release event was the same for both tau_P301L_ pos and tau_P301L_ neg (tau_P301L_ neg: 0.23 ± 0.01 μm, tau_P301L_ pos: 0.22 ± 0.01 μm; P = 0.91, two-tailed KS-test). This result shows that the increase in intensity in tau_P301L_ pos compared to tau_P301L_ neg is not due to an increase in MVR.

We next determined if the average pHluorin-VGLUT1 intensity per vesicle alone could explain observed increases in tau_P301L_ pos pHluorin-VGLUT1 intensity compared to tau_P301L_ neg. To determine single vesicle intensity, we utilized established integrated intensity of single release events (Maschi & Klyachko, 2017). Briefly, we used an integrated intensity box around single release events (Dotted Box, Fig. 5E), and plotted the integrated intensity as a function of time (Fig. 5F). We then quantified the difference between the background intensity and the average intensity after release (ΔF in Fig. 5F). Finally, we aggregated the quantized intensity (Fig. 5G) to compare tau_P301L_ pos and tau_P301L_ neg single vesicle release event pHluroin-VGLUT1 intensity.

For aggregated individual release intensities, the cumulative fraction of tau_P301L_ pos (Red Squares, Fig. 5G) integrated intensity distribution is statistically larger (P = 0.0046, two-tailed KS-test) than that of the tau_P301L_ neg condition (Black Circles, Fig. 5G). We then fit both distributions to a cumulative fit function (Blue Lines, Fig. 5G), and compared the resulting fit means to quantitatively determine the difference between tau_P301L_ pos and tau_P301L_ neg. We found the mean intensity for tau_P301L_ pos neurons was ∼1.4x times the tau_P301L_ neg intensity (tau_P301L_ neg: 1.09 ± 0.035, tau_P301L_ pos: 1.42 ± 0.07; P = 0.0046, two-tailed KS-test). This result supports the hypothesis that the increase in pHluorin-VGLUT1 intensity in tau_P301L_ pos compared to tau_P301L_ neg is due exclusively to increased single vesicle pHluorin-VGLUT1 intensity. Further, since pHluorin-VGLUT1 intensity comes from a single fluorophore per VGLUT1 transport, the increase in single vesicle pHluorin-VGLUT1 intensity is caused by an increase in the number of VGLUT1 transporters per vesicle, which we address in the discussion section.

### 2.6 Tau_P301L_ positive neurons exhibit increased VGLUT1 expression compared to tau_P301L_ negative neurons

To establish if tau_P301L_ pos exhibit increased VGLUT1 expression at 14-18DIV cultures, we used immunostaining of synaptophysin and VGLUT1 in cultured neurons (Fig. 6) from tau_P301L_ pos and tau_P301L_ neg littermates. We then imaged puncta of synaptophysin intensity (Fig. 6A), VGLUT1 puncta intensity (Fig. 6B), and accepted only co-localized puncta peaks to quantify relative puncta intensity (Fig. 6C).

**Figure 6:**
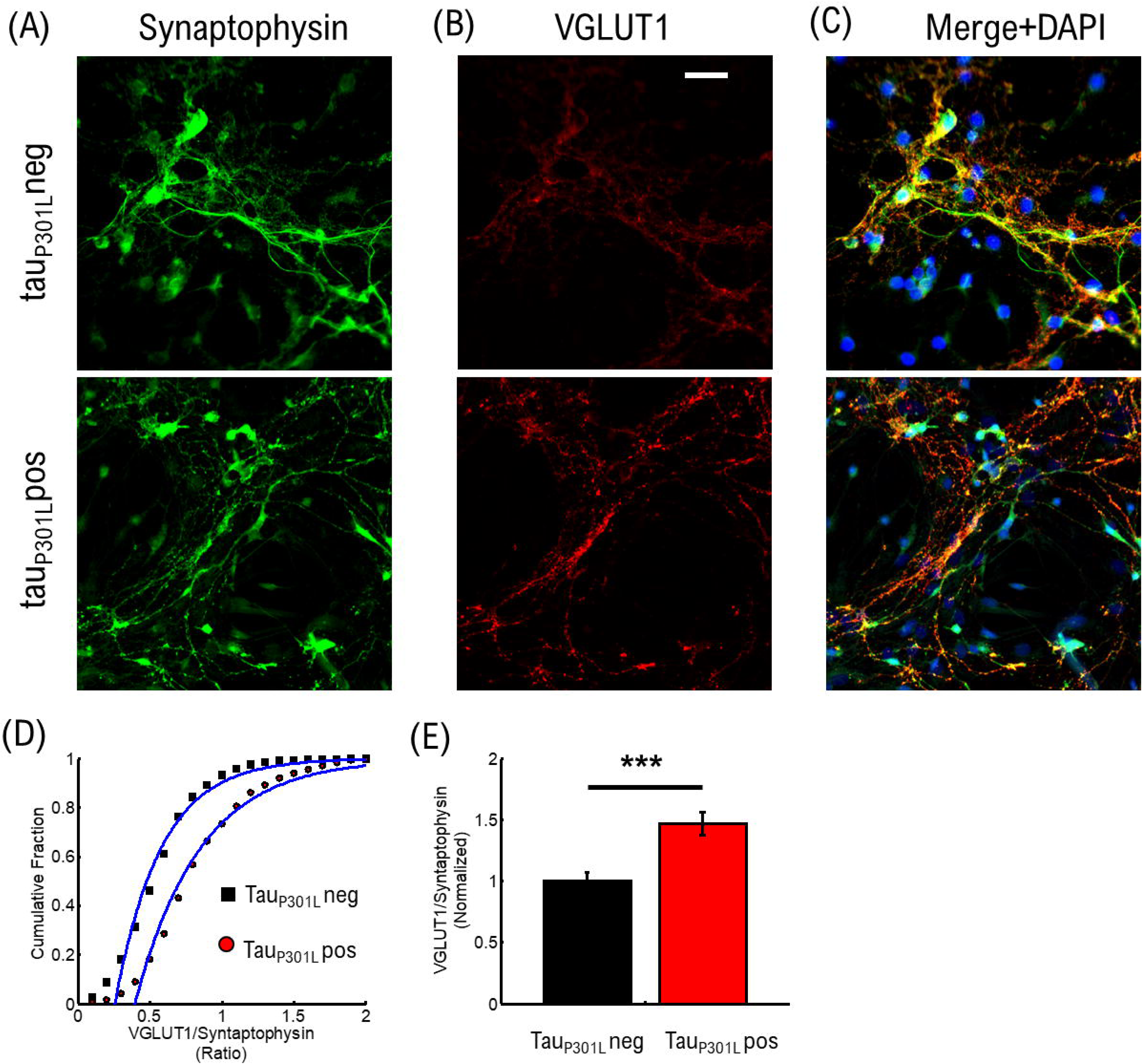
Tau_P301L_ pos neurons exhibit increased overall VGLUT1 expression compared to tau_P301L_ neg neurons. (A) Stained Synaptophysin intensity for tau_P301L_ neg (top), tau_P301L_ pos (bottom) (B) Stained VGLUT1 intensity for tau_P301L_ neg (top), tau_P301L_ pos (bottom) (C) Merged Synaptophysin (Green), VGLUT1 (Red), and DAPI (Blue) for tau_P301L_ neg (top), tau_P301L_ pos (bottom) (D) Cumulative distribution of VGLUT1/Synaptophysin intensity ratio for tau_P301L_ neg (black) and tau_P301L_ pos (red) (E) Mean ratio of VGLUT1/Synaptophysin intensity from fits in (D) and normalized to tau_P301L_ neg. Error-bars are errors of fit to data in (D). *** = P<0.001, two-tailed student t-Test.

We compared the ratio of VGLUT1/synaptophysin intensities to determine if an increase in VGLUT1 is observed. The cumulative distribution of intensity ratios showed a clear increase in VGLUT1 relative to synaptophysin for tau_P301L_ pos neurons compared to tau_P301L_ neg littermates (Fig. 1D). To determine the relative increase in VGLUT1, we then compared the fit mean values (Blue Lines Fig. 1D) and observed a relative increase in VGLUT1 of ∼1.48x (tau_P301L_ neg: 0.57 ± 0.07; tau_P301L_ pos: 0.85 ± 0.09; P < 0.001, two-tailed t-Test) in tau_P301L_ pos neurons (Fig. 6E). We note that the synaptophysin intensity distribution varied less than 4% between samples of the same litter and across litters for both tau_P301L_ pos neurons compared to tau_P301L_ neg (Supplemental Materials).

These results show that VGLUT1 expression is increased in tau_P301L_ pos mice compared to tau_P301L_ neg mice in cultured hippocampal neurons at 14-18DIV. Further, the 40-50% increase in VGLUT1 observed here is consistent with the 40% increase in VGLUT1 levels previously observed in adult tau_P301L_ pos mice (Hunsberger et al., 2014, 2015).

### 2.7 Tau_P301L_ pos neurons exhibit increased glutamate release as compared to tau_P301L_ neg neurons

The concentration of glutamate per vesicle has been shown to be proportional to the number of VGLUT1 transporters on the vesicle (Wilson et al., 2005). Increased glutamate concentration per vesicle would, in turn, result in an increased glutamate concentration in the synaptic cleft upon release. To establish the consequences of increased VGLUT1 transporters per vesicle in tau_P301L_ pos neurons, we measured extracellular glutamate release using the established glutamate reporter iGluSnFR (Marvin et al., 2018). The reporter is a membrane bound GFP fluorescent protein that becomes fluorescent when glutamate is released due to exocytosis and then bound to the reporter (Fig. 7A). Thus, the intensity of this reporter is proportional to the amount of glutamate released during synaptic transmission (Marvin et al., 2018).

**Figure 7:**
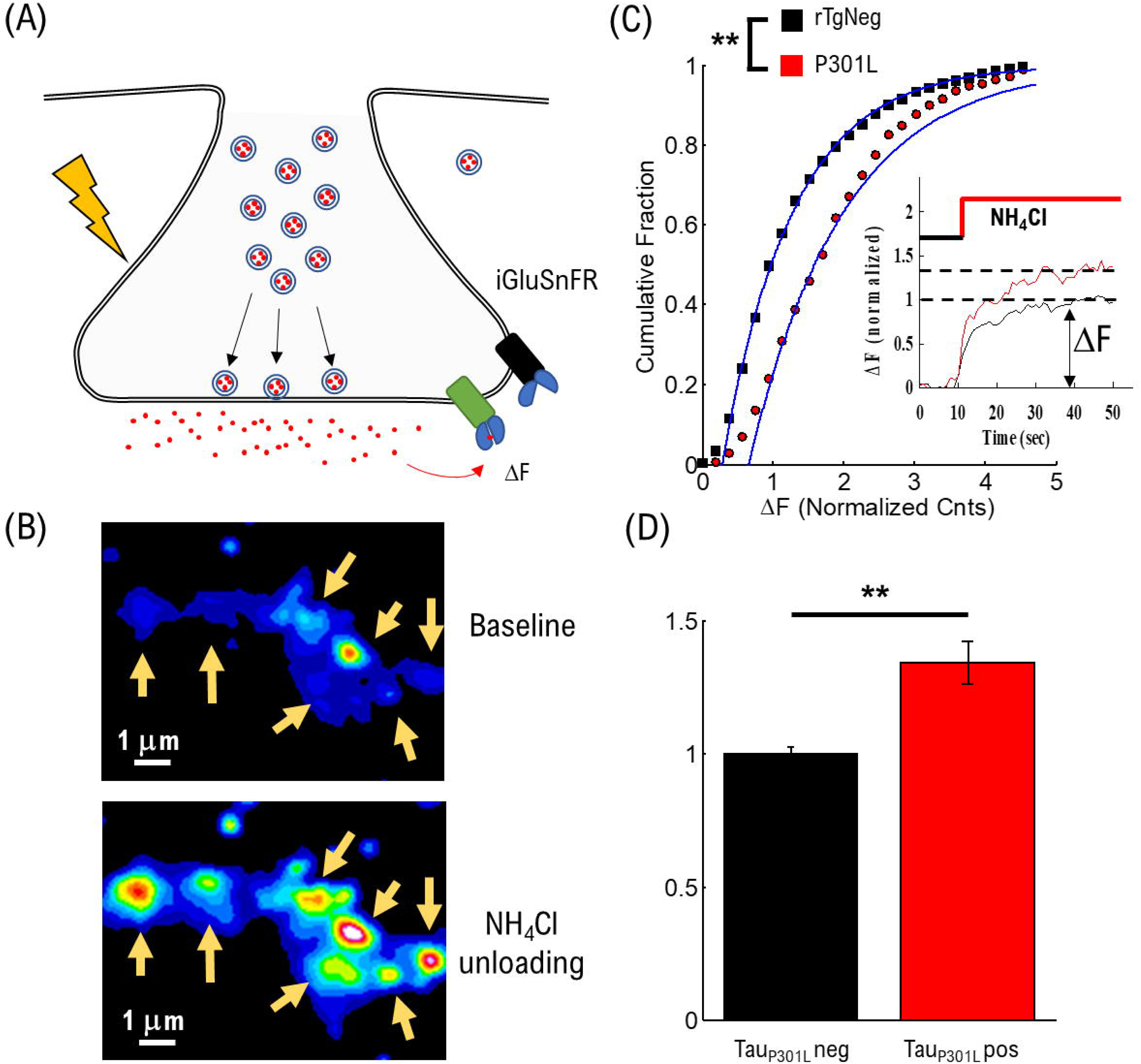
Tau_P301L_ pos exhibit increased overall iGluSnFR intensity compared to tau_P301L_ neg controls. (A) Representation of iGluSnFR fluorescence observed during presynaptic transmission (B) Example of raw iGluSnFR data observed before and directly after presynapse unloading induced by NH_4_Cl (C) Quantitative comparison of NH_4_Cl induced iGluSnFR intensity. Inset shows example traces during exposure to NH_4_Cl. (D) Comparison of mean iGluSnFR intensity increase for tau_P301L_ neg (Black) and tau_P301L_ pos (red). tauP301L neg N = 375 presynapses; P301L N = 526 presynapses; ** = P < 0.01 for 2-tailed t-Test. Error-bars are error of fits.

To determine the effects of increased VGLUT1 on overall extracellular glutamate release per synapse, we applied NH4Cl in the bath solution during imaging to depolarize the cellular membrane, which drives presynapses to unload the majority of their available vesicle pool (Fig. 7B) (Chanaday & Kavalali, 2018; Ganguly et al., 2015). Intensity level is low prior to addition of NH4Cl (Top image, Fig. 7B), and increases after addition of NH4Cl (Bottom image, Fig. 7B). Importantly, the overall spatial scale of intensity increase is the same as observed in pHluorin-VGLUT1 intensity (Fig. 1B), which supports that our glutamate measurement is localized to the presynaptic regions, consistent with previous iGluSnFR studies in hippocampal cell cultures (Marvin et al., 2018). We quantitatively compared intensity increase for iGluSnFR over the same spatial range surrounding identified presynapse locations as used in pHluorin to proportionally compare any changes.

We observed a significant increase in iGluSnFR intensity in tau_P301L_ pos as compared to tau_P301L_ neg controls (Fig. 7C). We compared aggregate cumulative fractions of increased fluorescence intensity after exposure to NH4Cl, and normalized to the mean tau_P301L_ neg intensity. Mean tau_P301L_ pos intensity increased by ∼1.4x compared to tau_P301L_ neg controls (tau_P301L_ pos: 1.4 ± 0.08; tau_P301L_ neg: 1.0 ± 0.03; P = 0.0025, two-tailed t-Test). This increased iGluSnFR intensity is proportional to both the overall increase in pHluorin intensity during stimulation (Fig. 1, 3) and increased single vesicle intensity (Fig. 7).

The iGluSnFR results support the hypothesis that increased VGLUT1 transporters per vesicle result in an increased extracellular glutamate release into the synaptic cleft.

## 3 Conclusions

In this present study we found tau_P301L_ pos mice exhibit increased VGLUT1 pHluorin intensity compared to tau_P301L_ neg littermates using the pH-sensitive fluorescent protein pHluorin-VGLUT1 (Fig. 1). We hypothesized that there were several pathways that could result in increased tau_P301L_ pos pHluorin intensity (Fig. 2). We found that the increased pHluorin intensity was not stimulation frequency dependent (Fig. 3). We further found that the release probability was the same for both tau_P301L_ pos and tau_P301L_ neg (Fig. 4). We then observed that the only single vesicles released for each stimulation pulse (Fig. 5) and that each single release event exhibited an ∼1.4x pHluorin-VGLUT1 intensity for tau_P301L_ pos as compared to tau_P301L_ neg (Fig. 5). We then found that tau_P301L_ pos mice exhibit increased VGLUT1 levels compared to tau_P301L_ neg littermates at 14-18 DIV (Fig. 6). Finally, we measured extracellular glutamate released using the fluorescent reporter iGluSnFR, and found that tau_P301L_ pos neurons exhibit a 1.4x increase in glutamate released as compared to tau_P301L_ neg littermates (Fig. 7). We conclude that increased single vesicle pHluorin intensity is caused by an increase in VGLUT1 per vesicle, and the increased VGLUT1 results in increased extracellular glutamate released during synaptic transmission.

## 4 Discussion

The results from this study provide key insights into presynaptic changes that may support observed cognitive changes during disease progression. The results here show that the number of vesicles released during synaptic transmission do not change in tau_P301L_ pos neurons, but the number of VGLUT1 transporters is increased. We can now use these results to estimate how many VGLUT1 transporters per vesicle exist in tau_P301L_ pos and how many vesicles are released during synaptic transmission, which can then be used to better model changes that occur later in disease progression.

### 4.1 Methodological inconsistencies of measurements of tauopathy mediated effects on presynaptic transmission

One major problem in understanding how tau mediates presynaptic pathological changes during disease progression is inconsistent results from different methodological approaches. Further, altered VGLUT1 levels and tau mislocalization in the presynapse can have confounding effects on presynaptic transmission. Therefore, it is essential to put the results from this study in the broader context of previous studies on tauopathy effects on presynaptic transmission.

(i) Our cellular culture approach differs from other cell culture approaches in keyways. We studied tau_P301L_ pos neurons grown on tau_P301L_ pos astrocytes and compared their differences to tau_P301L_ neg neurons grown on tau_P301L_ neg astrocytes. Previous studies have used immortalized hippocampal cells or neuroblastoma (SHSY5Y) cell lines to study tau-induced VGLUT1 levels, which have not been shown to be directly related to the time-course of tauopathy in P301L (Siano et al., 2019). Alternatively, primary hippocampal neurons in htau mouse lines have been grown directly on treated glass coverslips (McInnes et al., 2018; Zhou et al., 2017), but their results are physiologically limited because previous studies have also shown that neuron-astrocyte interactions can be a contributing factor in cell cultures of tauopathy models (Sidoryk-Wegrzynowicz et al., 2017). Further, cultured hippocampal neuron studies used transfection with a htau mutation to show a significant reduction in presynaptic transmission (McInnes et al., 2018; Siano et al., 2019; Zhou et al., 2017); however, it has not been shown how acute introduction of tau is directly related to the normal time-course of neurodegeneration. Therefore, our approach provides the advantage of measuring tau-mediated presynaptic vesicle release changes during their development, and in the presence of their appropriate astrocytic growth factors.
(ii) Our observation of VGLUT1 and our isolation of presynaptic transmission from post-synaptic signaling allows for a direct measurement of the contribution of VGLUT1 in presynaptic transmission in tauopathy models. Previous age-dependent *in vivo* mouse model studies have been able to correlate changes in VGLUT1 expression levels and their effect on calcium (Wu et al., 2021) or glutamate (Hunsberger et al., 2014, 2015), but these studies could not directly measure if VGLUT1 levels mediated increased extracellular glutamate; this is particularly important because changes in VGLUT1 levels have been shown to directly affect presynaptic release probability (Wilson et al., 2005, p. 1). Direct measurements of VGLUT1 levels on synaptic transmission using *in vitro* cell culture methods were inconsistent with *in vivo* results (Siano et al., 2019), possibly due to the limitations of the cell culture approach used, as outlined above. The approach in our study directly measures VGLUT1, extracellular glutamate, and blocks the post-synaptic response, thereby allowing for an isolated measurement on presynaptic transmission.
(iii) Our measurement of single vesicle VGLUT1 levels and single vesicle presynaptic release probability separates the confounding roles of VGLUT1 and vesicle release. It is not clear if presynaptic VGLUT1 levels increase prior to tau-mediated decreases in presynaptic transmission or if the two occur simultaneously. Previous *in vitro* studies of tau binding to single vesicles resulting in a reduction in presynaptic transmission did not measure VGLUT1 levels and were done using htau transfections (McInnes et al., 2018; Zhou et al., 2017). Alternatively, *in vitro* or *in vivo* cell culture VGLUT1 expression level experiments showed a change in extracellular glutamate levels but did not directly measure single vesicle presynaptic transmission (Hunsberger et al., 2015).

### 4.2 Number of VGLUT1 transporters per vesicle

Since single vesicle pHluorin-VGLUT1 intensity is caused by the intensity per VGLUT1 transporter, it is not physically possible for tau_P301L_ pos single vesicles to have 1.4x the number of VLGUT1 transporters as they must be whole integers. We can then use the single vesicle intensity distributions to estimate the number of VGLUT1 transporters per vesicle in both tau_P301L_ neg and tau_P301L_ pos. We first consider that the integer number of VGLUT1 transporters per vesicle in tau_P301L_ neg (N, Fig. 8A) results in the mean single vesicle intensity observed, and the distribution of number of VGLUT1 transporters equals the variance in observed intensity (σ, Fig. 8A). The observed increase in the tau_P301L_ pos gaussian distribution of intensity must then be constrained to an integer number (1.4*N, Fig. 8B), with an integer variance in the number per vesicle (σ, Fig. 8B). These constraints leave a limited number of potential options for the number of VGLUT1 channels that satisfy both the tau_P301L_ neg and tau_P301L_ pos distributions (Table 1).

**Figure 8:**
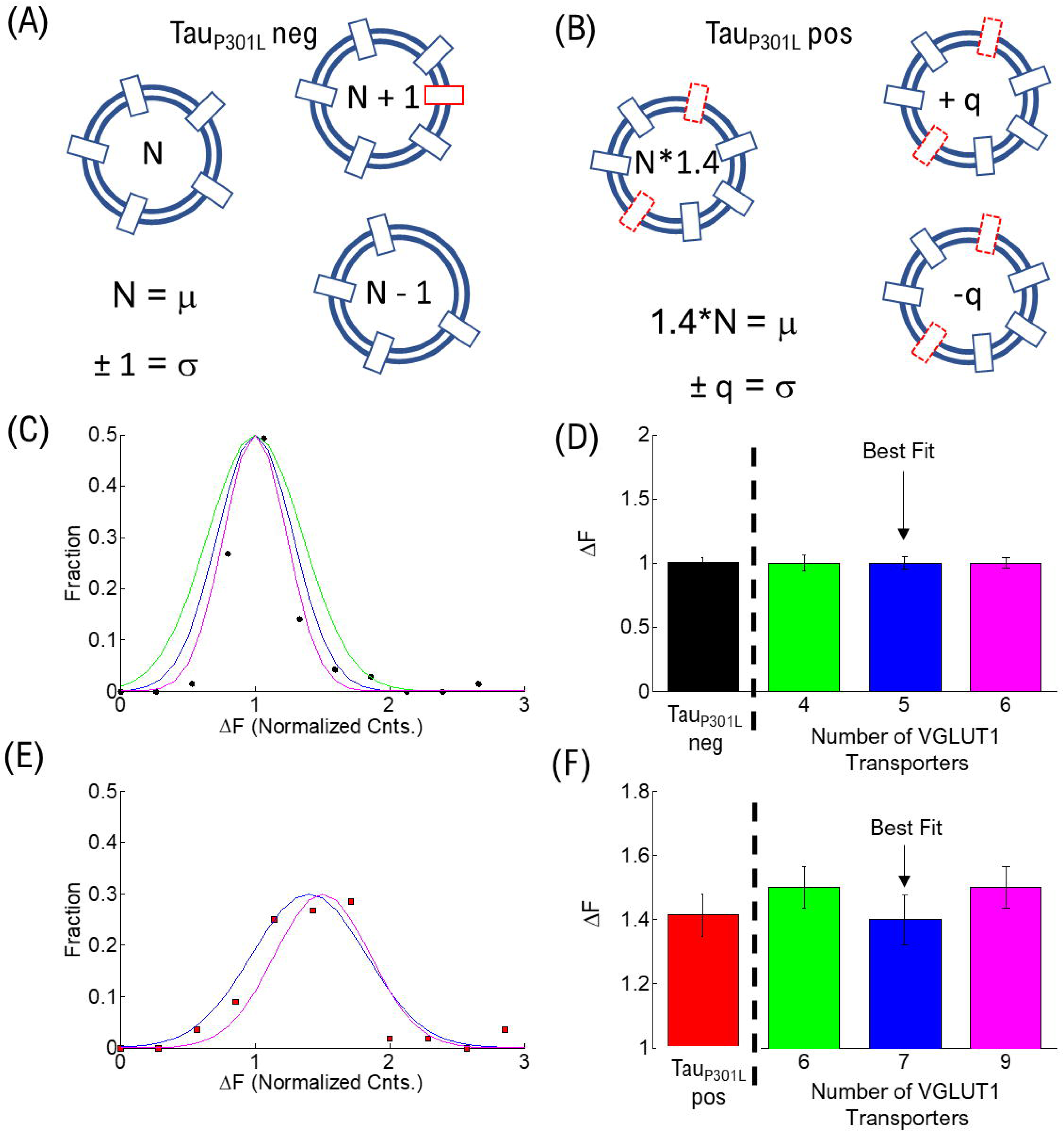
Estimate of Number of VGLUT1 transporters per vesicle. (A) Cartoon model of the number of VGLUT1 transporters per vesicle (N) and the variance in number (σ) (B) Cartoon model of the increase in number of VGLUT1 transporters in tau_P301L_ pos (N*1.4) and the variance in number (σ). (C) Fraction of single vesicle intensities per vesicle in tau_P301L_ neg (Solid circles) and modeled integer number of VGLUT1 transporters per vesicle of 4 (Green), 5 (Blue), and 6 (Magenta), with each having a variance of 1 per vesicle. (D) Normalized intensity per vesicle and variance for all modeled options of tau_P301L_ neg (Colors same as C). (E) Fraction of single vesicle intensities per vesicle in P301L (Solid squares) and modeled integer number of VGLUT1 transporters per vesicle equal to the nearest integer increase above tau_P301L_ neg (6 = 1.5*4, Green), (7 = 1.4*5, Blue), and (9 = 1.5*6, Magenta), with each having a variance of 1 per vesicle. (F) Normalized intensity per vesicle and variance for all modeled options of tau_P301L_ pos (Colors same as E).

**Table 1.**
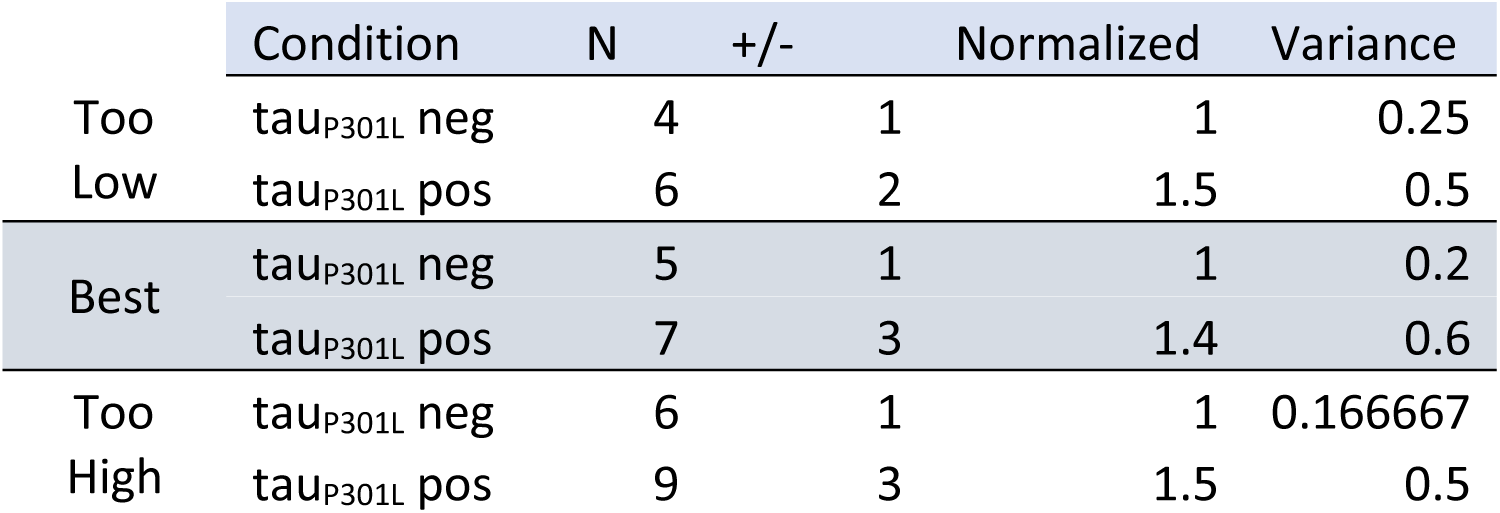

|  | Condition | N | +/- | Normalized | Variance |
| --- | --- | --- | --- | --- | --- |
| Too Low | $\text{tau}_{P301L}$ neg | 4 | 1 | 1 | 0.25 |
| | $\text{tau}_{P301L}$ pos | 6 | 2 | 1.5 | 0.5 |
| Best | $\text{tau}_{P301L}$ neg | 5 | 1 | 1 | 0.2 |
| | $\text{tau}_{P301L}$ pos | 7 | 3 | 1.4 | 0.6 |
| Too High | $\text{tau}_{P301L}$ neg | 6 | 1 | 1 | 0.166667 |
| | $\text{tau}_{P301L}$ pos | 9 | 3 | 1.5 | 0.5 |

We first modeled the tau_P301L_ neg single vesicle condition to find the minimum number of VGLUT1 transporters that would satisfy the observed result (Fig. 8 C,D). We found that a minimum of 4 transporters with a variance of 1 and a maximum of 6 transporters with a variance of 1 satisfied the constraints observed experimentally; below or above those values lead to values outside previously physically observed limits of VGLUT1 transporters (between 5 – 12 transporters per vesicle) (Takamori et al., 2006).

We then multiplied the number of VGLUT1 transporters per tau_P301L_ neg vesicle (4, 5, 6) by the factor that resulted in the next nearest integer number of vesicle per transporter; for example, an N = 4 transporters for tau_P301L_ neg would be multiplied by 1.5* to get an integer increase to 6 transporters per tau_P301L_ pos vesicle. We then found that for tau_P301L_ neg VGLUT1 to have both N = 4, 6 would require a factor of 1.5 times to get the nearest integer of VGLUT1 of 6 (Green, Fig. 8E,F) and 9 (Magenta, Fig. 8E,F) respectively in tau_P301L_ pos. The best fit result was the model of N = 5 VGLUT1 transporters for tau_P301L_ neg and N = 7 = 1.4*5 VGLUT1 transporters for tau_P301L_ pos (Blue, Fig. 8E,F).

### 4.3 Number of Vesicles Released During Stimulation

Aside from the main conclusion that our results show an increase in extracellular glutamate release from an increase in VGLUT1 expression per vesicle, we can now turn to focus on the question of how many vesicles are involved during sustained synaptic transmission. This question is important for two reasons: (1) the amount of extracellular glutamate is intimately tied to the number of vesicles released, and (2) any pathway that mediates neurodegeneration in AD using the presynaptic recycling pathway would be sensitive to the number of vesicles utilized during transmission and recycling. For example, the number of vesicles in the recycling pathway, which are predominantly used for synaptic transmission, is a relatively small fraction (∼20%) of all vesicles (Alabi & Tsien, 2012; Gross & von Gersdorff, 2016), and reserve pool vesicles are not predominantly used. Obtaining a quantitative comparison in the number of vesicles involved in transmission in tau_P301L_ pos neurons and tau_P301L_ neg littermates would thus provide an important measure on the potential number of vesicles, and which pool they may come from (Fig. 9A).

**Figure 9:**
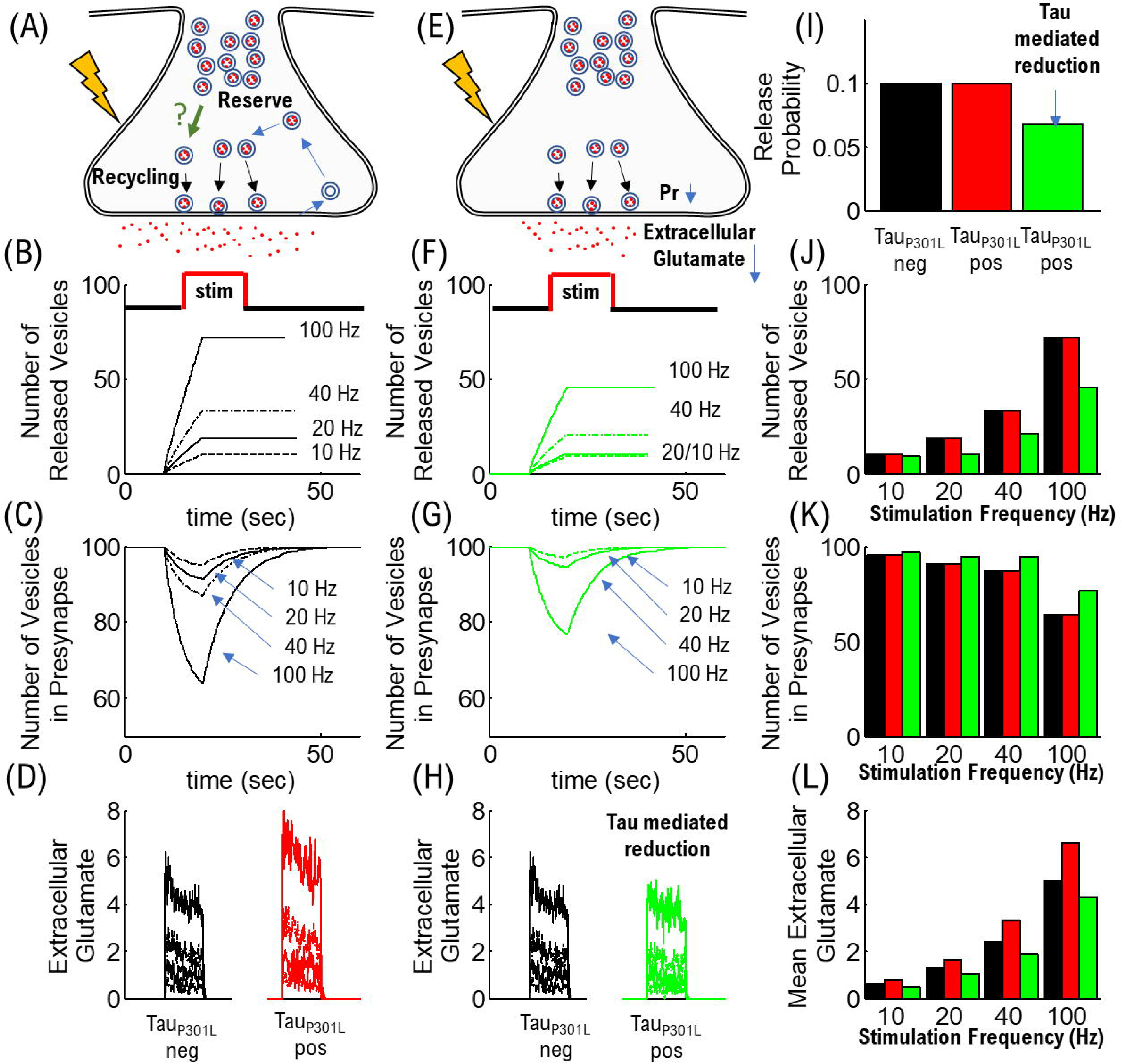
Estimate of Number of Vesicles Released During Stimulation. (A) Cartoon model of the number of vesicles utilized during sustained synaptic transmission (B) Number of vesicles remaining in the presynapse during stimulation (C) Number of unique release events during stimulation (D) Amount of extracellular glutamate released, normalized to 10 Hz. (E) Cartoon model of required reduced release probability to obtain hypothesized reduced extracellular glutamate (F) Number of unique release events during stimulation. (G)) Number of vesicles remaining in the presynapse during stimulation (H) Amount of extracellular glutamate released, normalized to 10 Hz. (I) Modeled reduction in release probability per vesicle (J) Number of unique release events during stimulation (K) Number of vesicles remaining in the presynapse during stimulation (L) Peak Amount of extracellular glutamate released, normalized to 10 Hz.

To address this important question, we use the computational model of presynaptic release probability (Fig. 3) utilized in our present study. We can separate the question of the number of vesicles used into two parameters: (i) how many vesicles remain in the presynapse during sustained synaptic transmission, and (ii) how many total release events are observed. The advantage of separating the question this way is that if only recycled vesicles are utilized during transmission as previously proposed (Alabi & Tsien, 2012), then any vesicle in the reserve pool is unlikely to contribute to changes observed.

We first look at the total number of release events to determine how many release events occur during a single bout of synaptic transmission. To determine total release, we simply counted each time a release event occurred during stimulation regardless of the number of times a vesicle may have already gone through the recycling process. We then compared the number of release events for different stimulation frequencies (Fig. 9B). The number of released vesicles increases with increasing stimulation frequency, as expected, however, because the probability of release is the same for both tau_P301L_ neg and tau_P301L_ pos synapses, the number of vesicles used is the same.

We next look at the total number of vesicles available in the presynapse during a single bout of stimulation. We assume the presynapse has a starting pool of 100 vesicles, that an exocytosis event removes a single vesicle from this pool, the removed vesicle is recycled and added back to the pool after a fixed endocytosis time. We use the same parameters for release probability and endocytosis rate as used in the results (section 2.3). With these assumptions, we compare the number of vesicles in the pool for different stimulation frequencies (Fig. 9C). The simulations show that the number of vesicles removed from the pool remain less than 20% for most stimulation frequencies up to 40 Hz, and a high stimulation frequency of 100 Hz is required to remove more than 20%. This suggests that the number of vesicles utilized during synaptic transmission remains consistent with the fraction of vesicles in the recycling pool for both tau_P301L_ pos neurons and tau_P301L_ neg littermates. Further, this suggests that increased vesicle release for stimulation frequencies greater than 40 Hz would require vesicles in the reserve pool and thus be sensitive to the mechanics of vesicles in the reserve pool.

Taken together, these results suggest that alterations in presynaptic transmission during AD progression are likely to be more sensitive to changes in the recycling vesicle pool than the reserve pool. However, during bouts of high stimulation (>40 Hz), as would occur during hyperexcitable states (Kazim et al., 2017), presynapses will begin to draw from the reserve vesicle pool.

### 4.4 Role of VGLUT1 transporters in extracellular glutamate release

Our results show that the amount of extracellular glutamate released increases, due to P301L tau mediated increase in VGLUT1 transporters per vesicle, even when the number of vesicles release is unchanged. Quantifying the specific number of VGLUT1 transporters per vesicle as well as the variance provides necessary information to understand the pathway to increased extracellular glutamate in the tau_P301L_ pos model. The specific number of VGLUT1 transporters is directly correlated with the amount of glutamate per vesicle (Wilson et al., 2005). Quantifying the number of VGLUT1 transporters in tau_P301L_ pos, as well as the variance in number, provides a constraint that can be used to better measure post-synaptic responses during the hyperexcitability state. This is particularly true when determining all the contributing factors of increased extracellular glutamate, such as glutamate re-uptake via astrocytic transporters (Anderson & Swanson, 2000). The relative contribution of extracellular glutamate from single vesicle release events can now be estimated based on the number of VGLUT1 transporters per vesicle. The extracellular glutamate per vesicle in tau_P301L_ pos is not simply 1.4x the normal amount, rather it is *specifically* resulting from 7± 3 VGLUT1 transporters per vesicle. However, the consequences of increased glutamate released per vesicle need to be understood in the context of an aggregate of release events.

To better understand the consequences of increased extracellular glutamate per vesicle, we used our computational model to estimate the increase in glutamate released as a function of stimulation frequency. To model the increased extracellular glutamate, we used our same computational model to estimate the number of vesicles released and used it to also model how much extra cellular glutamate is released (Fig. 9D). Here we assumed that extracellular glutamate is cleared at a rate <=100 msec (the time-step of our simulations) (Clements John D. et al., 1992). Further, we assume each vesicle releases the same amount of glutamate, the upper limit measured of ∼10,000 glutamate molecules per vesicle for tau_P301L_ neg (Wang et al., 2019), and 14,000 = 1.4* 10,000 for tau_P301L_ pos.

The simulation results show that extracellular glutamate present during stimulation is significantly increased at higher stimulation frequency (> 40 Hz) in tau_P301L_ pos (Red curves, Fig. 9D) compared to tau_P301L_ neg (Black curves, Fig. 9D). First, tau_P301L_ pos glutamate is 40% greater than for tau_P301L_ neg, for all stimulation frequencies. However, the difference in the overall amount of extracellular glutamate molecules present is relatively low for tau_P301L_ pos at 10 Hz (∼900 molecules) compared to tau_P301L_ neg (∼700 molecules); whereas the amount of extracellular glutamate molecules is significantly greater at 40 Hz for tau_P301L_ pos (∼31,000 molecules) compared to tau_P301L_ neg (∼25,000 molecules). The increased glutamate release becomes even larger for tau_P301L_ pos compared to tau_P301L_ neg at 100 Hz stimulation frequency.

This result suggests that the tau-mediated increase in VGLUT1, and in turn extracellular glutamate release, may not significantly contribute to a change in synaptic transmission at low stimulation frequencies but rather becomes a significant contributing factor during higher stimulation frequencies (> 40 Hz), as would be observed during hyperexcitable states.

### 4.5 Reduction in extracellular release due to tau mediated reduction in single vesicle release probability

We finally put our results of increased extracellular glutamate in context with the established observed tau-mediated decrease in synaptic transmission. Multiple forms of hyperphosphorylated tau have been shown to reduce synaptic transmission by reducing presynaptic vesicle recycling via binding to single synaptic vesicles and reducing vesicle mobility (McInnes et al., 2018; Wu et al., 2021; Zhou et al., 2017). However, this leads to two competing mechanisms to the amount of extracellular glutamate released. This level of extracellular glutamate is important based on the hypothesis that observed reduction excitatory post synaptic currents (EPSCs) are proportional to a reduction in extracellular glutamate. To put our results in context with tau mediated reduction in synaptic transmission, we ask the following question: what must the release probability be in order to compensate the increased glutamate per vesicle, in order to get an observed reduction in extracellular glutamate?

To address this question, we used out computational model to simulate a reduction in extracellular glutamate released. We first assumed that the tau-mediated reduction in vesicle exocytosis is modeled as a reduction in single vesicle release probability (Pr, Fig. 9E). We then assumed the glutamate released per vesicle is the same as the tau_P301L_ pos vesicles. We finally assumed that the reduction in extracellular glutamate is 20% lower than for the tau_P301L_ neg condition, which is within the 20-50% observed release reduction (McInnes et al., 2018; Zhou et al., 2017). We then used the simulated results to determine the number of vesicles released (Fig. 9F), the number of vesicles in the presynapse during stimulation (Fig. 9G), and the amount of extracellular glutamate molecules during stimulation (Fig. 9H).

With these assumptions, we found that a simulated release probability of Pr=0.068 (Fig. 9I) reproduced a 20% reduction in extracellular glutamate molecules during stimulation (Fig. 9L). More importantly, the number of vesicle release events during stimulation is approximately half the number used for both tau_P301L_ neg and tau_P301L_ pos with the release probability observed in this present study (Black & Red curves respectively, Fig. 9 J). Further, the tau-mediated reduction in release probability results in an increase in the number of vesicles remaining in the presynapse during stimulation compared to tau_P301L_ neg and tau_P301L_ pos with the release probability observed in this present study (Black & Red curves respectively, Fig. 9 K).

These simulated results show that the increase in VGLUT1 transporters per vesicle increases the amount of glutamate released during periods of higher stimulation (> 40 Hz), and requires a significant reduction in release probability to obtain the observed tau-mediated reduction in synaptic transmission previously observed in (McInnes et al., 2018; Zhou et al., 2017). This means that any approach to understand tau-mediated reductions in presynaptic glutamate release must include the counteracting mechanism of increased glutamate released per vesicle in order to correctly model synaptic transmission during disease progression.

## Supporting information

Key Resources

List of Figures and Data

## 5 Acknowledgements

MWG and MR would like to acknowledge Auburn University internal grant funding for support for the results in this manuscript.

## 6. Author Contributions

MWG and MR performed experimental design and analysis. ET managed mouse colony breeding, and sample preparation. MWG, ET, MH, JP performed experiments. MWG, ET, MH, JP, and MR analyzed experimental results. MG performed computational analysis. MWG, ET, MH, JP, and MR contributed to writing.

## Key Resources Table

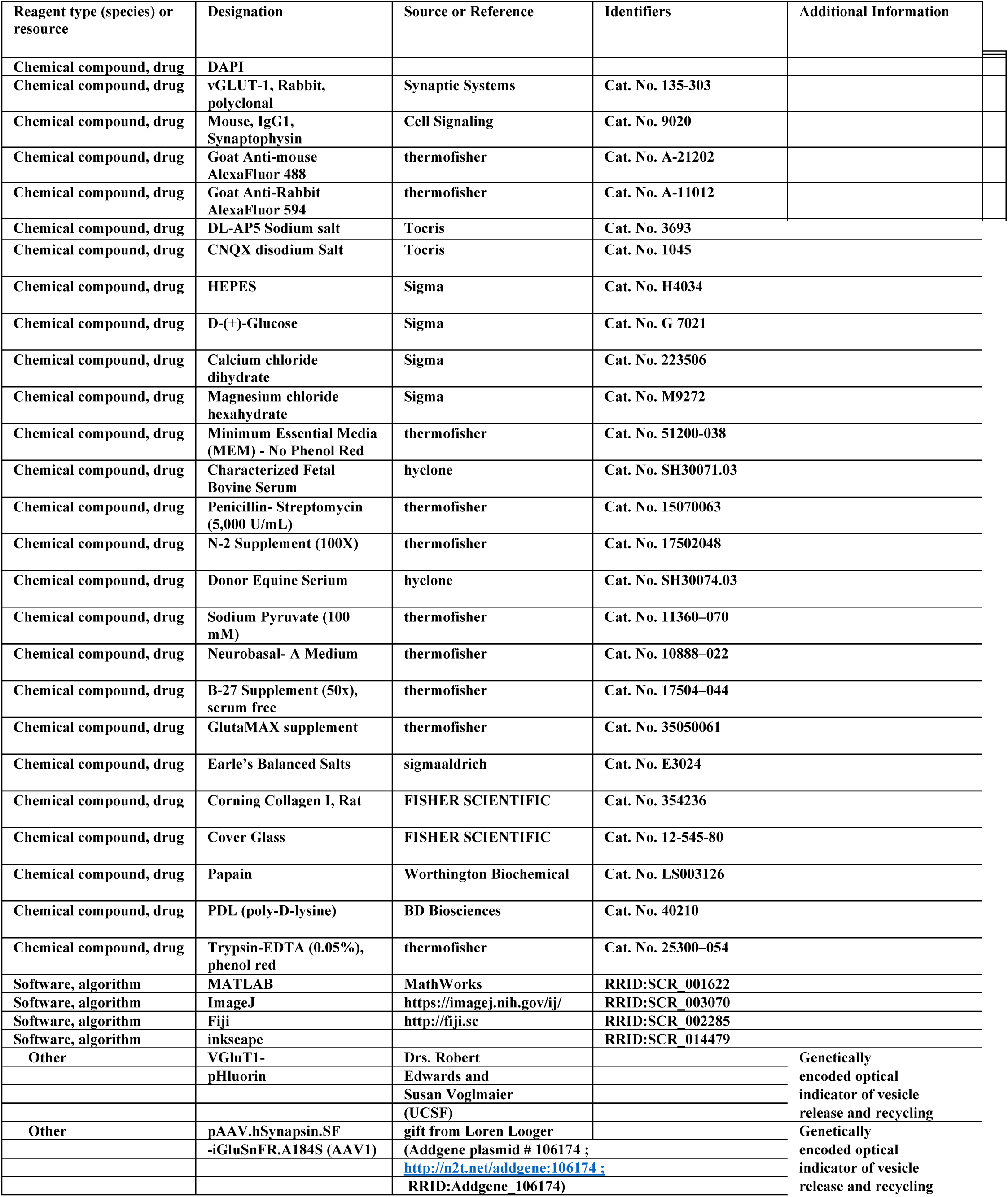

## Supplementary Material

### Animal Colony and Cell Culture Methods

TauP301L mice were created to represent cognitive decline, neurofibrillary tangle formation, and neuronal death that is seen in human Alzheimer’s Disease (Ramsden et al., 2005; SantaCruz K. et al., 2005). The TauP301L gene encodes for human four-repeat tau with P301L mutation (4R0N). Female TauP301L mice heterozygous for the tetracycline response element – TauP301L transgene were bred with male mice heterozygous for the activator gene: tet-off tetracycline transactivator downstream of the Ca^2+^/calmodulin kinase II promotor (CKtTA) (SantaCruz K. et al., 2005). The resulting mice were known as TauP301L/CKtTA. Based on PCR results, the resulting pups are designated +/+, -/-. +/-, or -/+. The first + or – denotes the presence or absence of the TauP301L pos gene. The second + or – denotes the presence or absence of the CKtTA activator gene. CKtTA activates TauP301L pos and creates the characteristics of Alzheimer’s Disease. Mice with both genes (+/+) were used to represent Alzheimer’s Disease and -/+, +/-, and -/- mice were used as negative controls. Since they do not have both the responder and the activator genes, the Alzheimer’s Disease phenotypes will not be displayed (Hoover et al., 2010; Ramsden et al., 2005; SantaCruz K. et al., 2005).

Around 16-20 TauP301L females and 4 CKtTA males are used for consistent rotation and bred following. Females were grouped into pairs and were housed together unless they were pregnant or with a litter. The day a litter is born is designated post-natal day 0 (PND 0). On PND 3 or 4, the pups receive paw tattoos and tail snips to use for PCR. On PND 4 or 5, their hippocampi are dissected, and the cells are cultured for experiments.

Cell cultures were plated on glass coverslips in 12-well plates made. One week prior to dissections two 12-well plates were plated with a uniform confluent layer of astrocytes. One plate contained only +/+ astrocytes (Plate 1, which we call tau_P301L_ pos). A second separate plate (Plate 2) contained +/-, -/+, -/- astrocytes combined (which we call tau_P301L_ neg). All animal dissections occurred on PND 4 or 5 and followed previously established protocols, and in accordance with approved IACUC protocols. All +/-, -/+, -/- pups were dissected, combined, and plated at the same time on Plate 2 (which we call tau_P301L_ neg). All +/+ pups were dissected separately and plated on the +/+ only astrocytes in Plate 1 (which we call tau_P301L_ pos). All neurons, regardless of condition, were plated at a density of 20k per plate. 12 hours after plating cell media was changed to a neurobasal medium for neuronal growth.

Plated cells were transfected following established pHluorine-VGLUT1 lentiviral vector methods(Maschi et al., 2018, 2021; Maschi & Klyachko, 2017; Voglmaier et al., 2006). Briefly, cells were transfected with 1mL of virus+Neurobasal medium with MOI ∼1 at 3DIV. Cell media was then changed to back to neurobasal growth medium after 48 hrs of virus exposure. Every 5 days half the media was replaced with fresh neurobasal media. Cells were maintained in a cell culture incubator before imaging between 13 - 18 days.

For iGluSnFR experiments, plated cells were transfected with the pAAV.hSynapsin.SF-iGluSnFR.A184S (Addgene) vector, following established AAV viral vector methods (Marvin et al., 2018). Briefly, cells were transfected with 1mL of virus+Neurobasal medium with MOI ∼40 at 3-4DIV. Cell media was then changed to back to neurobasal growth medium after 48 hrs of virus exposure. Every 5 days half the media was replaced with fresh neurobasal media.Cells were maintained in a cell culture incubator before imaging between 13 -18 days.

### Experimental Approach and Methods

#### Immunocytochemistry

Immunocytochemistry was performed similar to previously described (Glynn et al 2006). Briefly, media was removed from coverslips of healthy neurons were fixed with a solution of cold 4% paraformaldehyde in 0.1M phosphate buffered saline (PBS) with 4% sucrose for 10 minutes at 4°C. The fixative was quenched by washing with 0.1M glycine for 5 minutes and then washed three times. Cell membranes were permeabilized with 0.25% Triton X-100 for X min and samples were washed. Nonspecific binding was then blocked with incubation in 10% Normal Goat serum for 30 minutes. Primary antibodies were added and incubated in 1% Normal Goat serum in 0.1M PBS at 4°C overnight. Cells were then washed before secondary antibody incubation for 2 hours. Cells were washed and a DAPI counterstain was added for 5 minutes. Cells were washed one more time and mounted onto gelatin coated slides with Permount and fixed. All steps were performed at room temperature on a shaker at 40 rpm in the dark unless otherwise noted. All solutions were made using 0.1M PBS diluent. Samples were imaged on a Nikon Ti2 Fluorescence microscope.

#### Experimental Setup for pHluorin and iGluSnFR

Samples were imaged on a custom built microscope based on a Nikon Ti2-E inverted microscope base (Nikon), surrounded by a temperature controlled incubation chamber held at 37C (OKOLab). Samples were illuminated using an LED source (SOLA), with excitation and emission frequencies filtered using a GFP cube (Nikon). Intensity was focused onto samples using a 100x oil objective (Nikon). Emitted light intensity was collected using an Orca Flash v4 CMOS camera (Hamamatsu). Images were taken at 80 msec exposure followed by a 20 msec readout time. Images were recorded using Hamamatsu software in *.cxd format and converted to individual *.tif files later for analysis (Hamamatsu). Field stimulation was applied using a square-pulse stimulator (BKPrecision 4030 10 MHz generator), with 1msec pulse durations at 10V/cm. Samples were kept alive during experiments and perfused using a multi-channel perfusion system (World Precision Instruments) with a modified tyrodes solution pH-balanced to 7.4 following previously established protocols. {Dario} Timing and control of all equipment was performed using a Marter9 TTL controller (AMPI Instruments).

#### Bulk pHluorin Analysis

Bulk pHluorin intensity curves were obtained from aggregating individual presynapses using the same process previously used for dye loading/unloading (Gramlich & Klyachko, 2017; Maschi et al., 2021). Briefly, raw tiff files were background subtracted using a 30-pixel rolling ball radius in ImageJ; second, single presynapses were identified and integrated using a 15×15 pixel box in custom written Matlab code; third, each presynapse pHluorin curve was background subtracted so that all curves were zero intensity just before stimulation (10 sec, see Fig. 1); then, the counts for each presynapse was binned for all curves as a function of frame and fit to a gaussian curve; Finally, the resulting mean +/- SEM as a function of frame was plotted for each condition.

#### Single Vesicle pHluorin Analysis

Single pHluorin intensity release measurements in Fig. 3,4 were obtained from aggregating individual presynapse release events using similar protocols previously used for single vesicle pHluorin experiments (Maschi et al., 2018, 2021; Maschi & Klyachko, 2017). Briefly, we background subtracted raw data using a 10-pixel rolling ball radius in ImageJ; we then applied a 1-pixel Gaussian blur in ImageJ; a 15×15 pixel integration box was used to obtain intensity from each identified presynapse location; intensity was then run through an 8-frame moving-average filter similar with our previously established approach (Gramlich & Klyachko, 2017; Maschi et al., 2018, 2021).

#### Bulk iGluSnFR Analysis

Bulk iGluSnFR intensity curves were obtained from aggregating individual presynapses using the same process previously used for pHluorin intensity. Briefly, raw tiff files were background subtracted using a 30-pixel rolling ball radius in ImageJ; second, single presynapses were identified and integrated using a 15×15 pixel box in custom written Matlab code; third, each presynapse pHluorin curve was background subtracted so that all curves were zero intensity just before NH4Cl exposure (50 sec, see Fig. 7); then, the mean peak intensity was determined by averaging the 10 frames with the most intensity for each presynapse; Finally, the resulting mean +/- SEM as a function of frame was plotted for each condition.

### Computational Simulation Approach for Presynaptic Vesicle Release

Computational simulations were performed using a model based on the established binomial vesicle release model, as shown in eqns. {2}, {3}. All model parameters used to reproduce each experimental conditions were constrained (Table S1) using the tau_P301L_ neg 40 Hz data (Black circles Fig. 2 C). Only the intensity per vesicle (q) and release probability (p_0_) were allowed to change in order to match observed rTgPos 40 Hz condition (Red Squares Fig. 2 C). Further, we modeled an exponentially decaying vesicle release probability with time during stimulation (𝑝(𝑡) = 𝑝_0_ + Δ𝑝 ∗ 𝑒^−𝑡/𝜏_𝑝_^) which course-grains a combination of factors that have been extensively explored previously, but are beyond the scope of this present study. The same time-dependent reduction response was used for both tau_P301L_ neg and rTgPos conditions and reproduced the same intensity curves.

Simulations were performed following the same algorithm approach using python 3.10 release. All simulation parameters (Table S1) are fixed at the beginning of each simulation. A single array (**I**) is created with length equal to the total number of simulation time-steps. Each time-step (Δt) in our simulations were equivalent to a single frame exposure time (100 msec). Each array element (**I**(Δt)) stores the total observable intensity (background + vesicles) equivalent to counts observed experimentally. We used a dynamic Monte-Carlo approach, that we have previously used to model vesicle probabilities(Gramlich et al., 2021; Gramlich & Klyachko, 2017), to model each probabilistic process in simulations of vesicle release mechanics.

The algorithm steps for each simulation time-step were as follows:

[1] The simulation sets the initial background intensity at Δt = 0.
[2] When the simulation time-step equals the stimulation start-time (Δt = Stim start), an unweighted random number (R_1_) are chosen between 0 and 1.
[3] If R_1_ is below the current release probability (p(Δt)), then a single vesicle is released with intensity (q) which is added to the total intensity for the array (**I**(Δt) = **I**(Δt)+q).
[4] A random number (R_2_) is then chosen from an exponential random number generator (np.random.exponential) seeded with the endocytosis rate (τ_endo_). This number represents the number of simulation time-steps that the released vesicle in [3] is counted in the array. Array elements from the current time-step (**I**(Δt) += q) up to the randomly generated time-step (**I**(Δt + R_2_) += q) have the vesicle intensity added.
[5] The algorithm steps [2]-[4] are repeated at the same time-step (Δt) for up to the number of experimental pulses used (Pulse/frame).
[6] The total intensity for the current simulation time-step is reduced by the amount fixed by the photo-bleaching parameter (PR)
[7] Algorithm steps [2]-[6] are repeated until the simulation time-step equals the end of the stimulation (Δt = Stim end)
[8] After stimulation, the remaining simulation time-steps reduce the intensity of each element by the amount fixed by the photo-bleaching parameter (PR)

To reproduce averaged pHluorin-VGLUT1 intensity results, we simulated 100 presynapses and reported the averaged simulated intensity. For each hypothetically proposed pathway, (ii)-(iii) in section 2.2, we ran 100 simulations. We then averaged the resulting intensity for all 100 simulations, and compared the averaged intensity with experimental results (Fig. 2B).

### Statistical Analysis Methods

For single vesicle width analysis, we performed a power analysis in Matlab for a t-test statistic assuming a tau_P301L_ neg = 250 +/- 50 nm baseline distribution to determine the number of samples required to for a tau_P301L_ pos = 300 +/- 50 nm distribution. For a power of 0.99, the analysis recommended a minimum of 21 samples per distribution. Thus, both the tau_P301L_ neg and tau_P301L_ pos sample numbers were sufficient to distinguish the difference if it existed.

### Presynaptic density and size is the same in tau_P301L_ pos and tau_P301L_ neg

To determine if the density of presynapses changes in tau_P301L_ pos compared to tau_P301L_ neg, we used integrated intensity line analysis. We used pHluosin-VGLUT1 intensity data after samples were exposed to NH_4_Cl, causing a complete unloading of all presynapses (Ganguly et al., 2015). We used ImageJ to open single raw tiff files at the frame of peak intensity (Fig. S1 A). We drew a line along an axon starting at a presynapse and ending just after a third neighboring presynapse. We then normalized the pHluorin-VGLUT1 intensity to the first presynapse. Finally, we aggregated multiple lines to determine an average intensity as a function of distance from the first presynapse (Fig. S1 B). We found that the pHluorin-VGLUT1 intensity decreased as a function of distance and begin to increase again until reaching a peak the first nearest neighbor (1^st^ nearest neighbor, Fig. S1 B). The pHluorin-VGLUT1 intensity then began to decrease again, and then rise again peaking at the second nearest neighbor (2^nd^ nearest neighbor, Fig. S1 B). The nearest neighbor peaks occurred at approximately the same location for tau_P301L_ pos (2.6 μm) compared to tau_P301L_ neg (2.86 μm). We note that there was a slight shift in the tau_P301L_ pos distance between presynapses, as well as an increase in the distribution of distances, but the difference did not significantly alter the density of presynapses along the axon. Further, these distributions are consistent with our previously observed distributions for rat hippocampal cell cultures (Gramlich & Klyachko, 2017).

Separately, we sought to determine if the overall size of the presynapse changed in tau_P301L_ pos compared to tau_P301L_ neg, which would potentially alter the total amount of pHluorin-VGLUT1 intensity. To determine this possibility we used the same pHluorin-VGLUT1 line intensity integration above, but for single presynapses. We used ImageJ to open single raw tiff files at the frame of peak intensity (Fig. S1 C). We drew a line along an axon across a presynapse. We then normalized the pHluorin-VGLUT1 intensity to the first presynapse. Finally, we aggregated multiple lines to determine an average pHluorin-VGLUT1 intensity across a presynapse (Fig. S1 D). We found that the pHluorin-VGLUT1 intensity FWHM (2σ, Fig. S1 D) was slightly smaller for tau_P301L_ pos (0.76 μm) compared to tau_P301L_ neg (0.82 μm). However, these differences are not significant compared to the uncertainty in the data, nor would the difference significantly contribute to the observed differences in overall pHluorin-VGLUT1 intensity observed.

### Variance in Synaptophysin expression measured in tau_P301L_ pos and tau_P301L_ neg

To determine if the synaptophysin distribution changed in immunohistochemistry analysis of tau_P301L_ pos compared to tau_P301L_ neg, we compared the distribution of intensities between samples within a litter and across litters. Immunohistochemistry data included 2-3 plates for each litter. We aggregated the distribution for all plates in a litter and used the width of the distribution (σ) as the variance between samples in a litter (Fig. S2 A,C). We then compared the SEM across three litters used for each condition to determine changes between litters (Fig. S2 B,D). We found that all litters had a variance less than 4% for both tau_P301L_ pos and tau_P301L_ neg.

---

**Figure S1:**
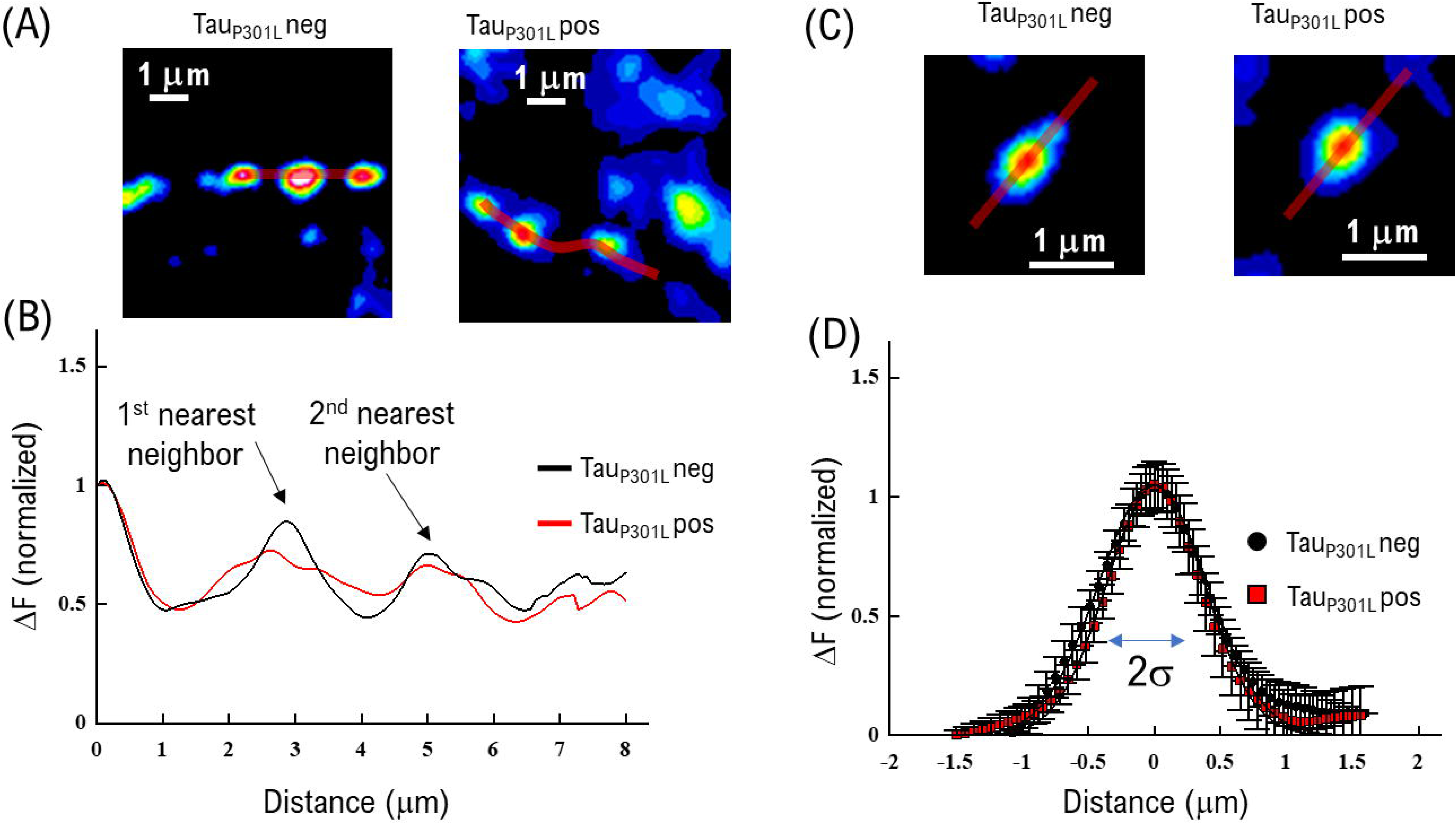
Estimate of density of presynapses and presynapse size. (A) Example of intensity integration line used to determine distance between presynapses (B) Aggregated integrated pHluorin-VGLUT1 intensity (Red line indicated in (A)) for tau_P301L_ pos (Red, N = 12) and tau_P301L_ neg (Black, N = 10). Nearest neighbor peaks are indicated. (C) Example of intensity integration line used to determine width of presynapses (D) Aggregated integrated pHluorin-VGLUT1 line intensity of individual presynapses for tau_P301L_ pos (Red, N = 12) and tau_P301L_ neg (Black, N = 10).

**Figure S2:**
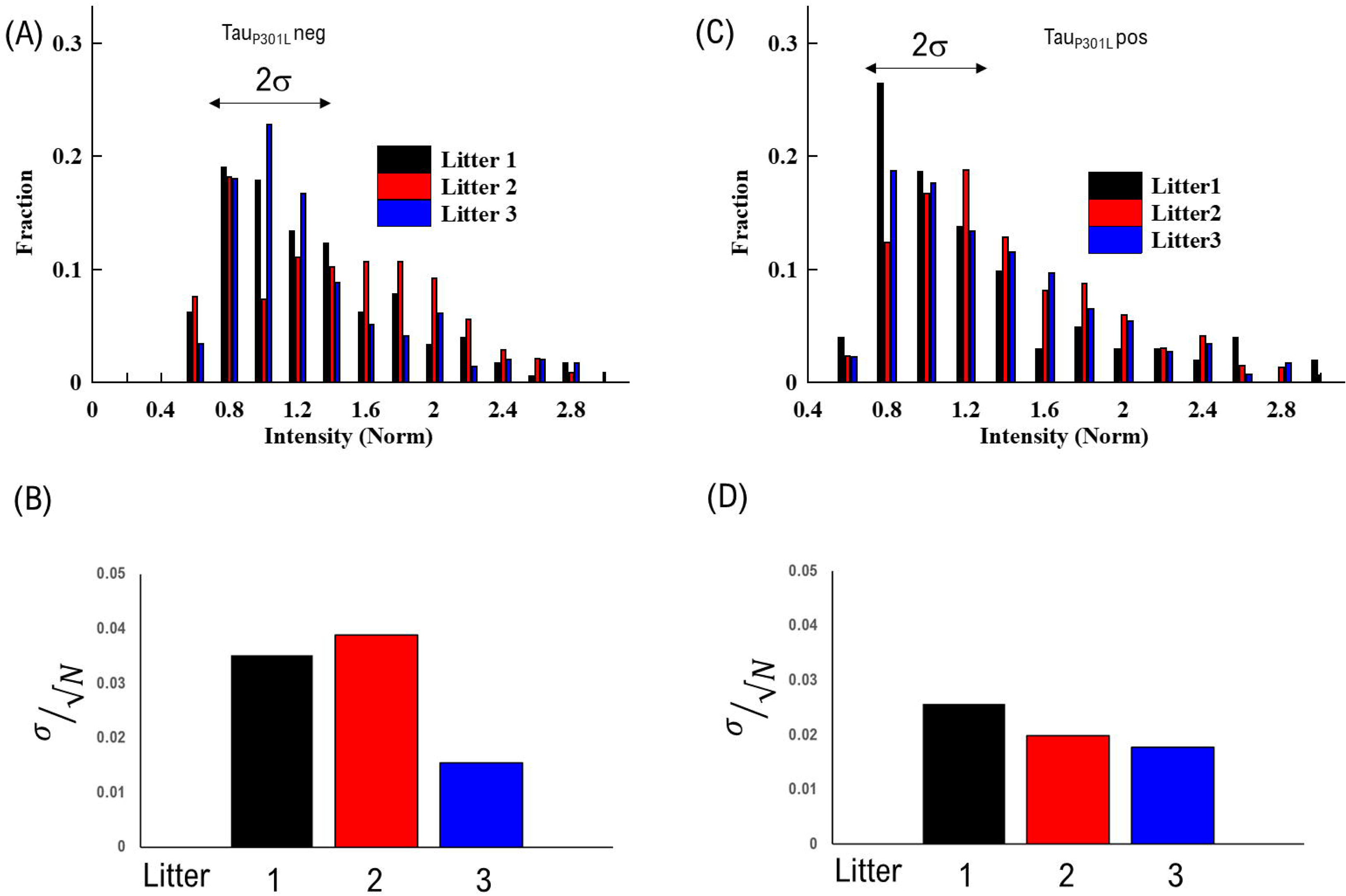
Variance in Synaptophysin measured with immunohistochemistry. (A) Distribution of Synaptophysin intensities normalized to mean across three litters measured for tau_P301L_ neg (B) SEM measured for distribution of synaptophysin intensities for tau_P301L_ neg (C) Distribution of Synaptophysin intensities normalized to mean across three litters measured for tau_P301L_ pos (D) SEM measured for distribution of synaptophysin intensities for tau_P301L_ neg

**Table S1:** Computational Simulation Parameters

| Parameter | Value | Purpose |
| --- | --- | --- |
| Pulse/frame | 4 | Number of stimulation pulses per each frame |
| $\tau_p$ | 7 (sec) | Rate of reduction in release probability during stimulation |
| $\Delta p$ | 1.6 | Release probability pre-factor |
| Stim start | 10 (sec) | Time stimulation starts |
| Stim end | 20 (sec) | Time stimulation ends |
| $\tau_{\text{endo}}$ | 7 (sec) | Endocytosis Rate |
| BKG_start | 1000 (cnts) | Background Intensity at start of simulation |
| BKG_end | -2000 (cnts) | Background Intensity at end of simulation |
| $\tau_{\text{BKG}}$ | 10 (sec) | Rate of decay in background intensity from photobleaching |
| PR | 0.1 | Fraction of intensity lost due to photobleaching at each time-step |
| $p_0$ | free | Release probability |
| $q$ | free | Intensity per vesicle |

## Notes

### Competing Interest Statement

The authors have declared no competing interest.

